# FNIP1 Modulates B Cell Receptor Signaling Strength by Coordinating Metabolism During Development

**DOI:** 10.64898/2026.03.30.715168

**Authors:** Heon Park, Ryan Culbert, Dechen Sakya, Raynah R. Silprasert, Brian M. Iritani

## Abstract

B cell development relies on stringent checkpoints that ensure immune competence and eliminate autoreactive clones. Transitional B cells (B220⁺CD93⁺), which emerge from the bone marrow, migrate to the spleen and differentiate into follicular (FO) or marginal zone (MZ) B cells, a process governed by B cell receptor (BCR) signaling strength, metabolic fitness, and survival cues. Here, we identify Folliculin Interacting Protein 1 (Fnip1) as a key regulator of this developmental transition. Using conditional Fnip1-deficient mice (*Fnip1^fl/fl^CD21Cre*), loss of Fnip1 results in a developmental arrest at the transitional B220⁺CD93^mid^ stage, severely limiting differentiation into FO and MZ B cells and leading to accumulation of a distinct enlarged CD19^high^, RAG negative B cells. Fnip1 modulates BCR signaling thresholds and metabolic programming by regulating the AMPK/FLCN/TFEB and CD19/PI3K/Akt/mTORC1 pathways through restricting TFEB access to the nucleus. Using the MD4/mHEL/sHEL tolerance model, we show that Fnip1 is dispensable for negative selection but is essential for maintaining peripheral tolerance. Together, our findings define Fnip1 as a metabolic gatekeeper that integrates nutrient-sensing pathways with BCR signaling to orchestrate transitional B cell fate decisions, promote peripheral tolerance, and maintain immune homeostasis.

## INTRODUCTION

Mature B cells are replenished from transitional immature B cells that undergo stringent immune checkpoints to prevent autoreactivity and establish self-tolerance ^1^. These checkpoints, active in both bone marrow and spleen, are critical for ensuring that mature B cells respond to foreign antigens without targeting self-antigens. Immature B cells express surface IgM (slgM) formed by the pairing of Ig heavy (IgH) and light (IgL) chains. Upon encountering self-antigens, high-affinity autoreactive B cells are either deleted via negative selection or undergo receptor editing—reinitiating IgL chain rearrangement (κ or λ). B cells with low or no self-reactivity migrate to the spleen, where they continue to mature ^2–5^.

Transitional B cells, characterized by B220^+^CD93^+^ expression, progress through distinct developmental stages—T1, T2, and T3—marked by differential IgM and CD23 expression ^6^. These subsets undergo further selection before committing to either the follicular (FO) or marginal zone (MZ) B cell lineages. Transitional and FO B cells that evade deletion may enter an anergic state characterized by reduced B-cell receptor (BCR) surface expression and diminished responsiveness to self-reactive antigens ^4,7^.

BCR signaling strength is pivotal in directing transitional B cell fate. High affinity for self-antigens can lead to negative selection, while lower affinity, productive signaling promotes survival and maturation. The developmental transition from T1 to T2 to either FO or MZ B cells involves the fine-tuning of BCR signals in concert with inputs from receptors such as BAFF-R, Notch2, Toll-like receptors, integrins, and chemokines ^8–12^. The role of the BCR and associated signaling molecules in splenic B cell development has been extensively investigated using mouse genetic models. Studies of CD19, Foxo1, Notch2, and BAFF-R deficiency reveal diverse effects on transitional, MZ, and FO B cell maturation ^8,9,13–15^. Likewise, proximal BCR signaling components, such as Igα (mb1), Syk, BLNK, Btk, Lyn, CD45, Vav/Rac, PI3K-Btk-PLCγ2, and the metabolic signaling axis AKT/mTOR are essential for early transitional B cell progression and maturation by contributing to metabolic fitness and survival ^1,16,17^.

Recent work has comprehensively depicted the molecular mechanisms by which Folliculin (Flcn) and its binding partners, Folliculin-interacting protein 1 (Fnip1) and Fnip2, play a role in metabolic adaptation in the context of the AMPK/FLCN/Fnips/TFEB and mTORC1 axis ^18–20^. During nutrient shortage and lysosomal stress, activated AMP-activated protein kinase (AMPK) coordinates a cascade of signaling pathways to suppress anabolic pathways through the inhibition of FLCN/Fnip1 GAP activity, where mTORC1 promotes processes like protein, lipid, and organelle biosynthesis. In turn, this leads to the dissociation of RagC, mTORC1, and TFEB from the lysosome, resulting in the translocation of cytoplasmic TFEB to the nucleus. Active TFEB-mediated transcription of lysosomal genes results in mitochondrial biogenesis.

Previous work has shown that Fnip1-deficient mice and humans with Primary Immunodeficiency Disease due to loss-of-function mutations in *FNIP1* exhibit near complete blocks in B-cell development at the large pre-B stage, despite normal IL-7-mediated proliferation ^21–23^. These pre-B cells show heightened sensitivity to nutrient deprivation, and resistance to Myc-induced transformation. Fnip1-deficient T-cells are developmentally well preserved but invariant natural killer T (iNKT) cell development is largely impaired, resembling phenotypes observed in c-Myc and TSC1 deficiency ^24–26^.

In this study, we deleted Fnip1 specifically in mature B cells using the Cre-LoxP system to gain insight into the role of Fnip1 in mature B cell development and functions. The data reveal that Fnip1 serves as a metabolic gatekeeper that modulates BCR signaling strength, in part via the CD19/PI3K/AKT pathway, to adapt to metabolic demand.

## RESULTS

### Stage-specific Cre models reveal that Fnip1 functions as a gatekeeper regulating the progression of B cell development

Previous works revealed that Fnip1 deletion using whole body deletion (*Fnip1^-/-^)* or early pre-B cell specific deletion (*Fnip1^fl/fl^ Mb1Cre* mice) resulted in a complete block at the large pre-B cells ^21,23^. To examine if Fnip1 plays a role in B cell development beyond large pre-B cells, *Fnip1^fl/fl^* mice were crossed to stage specific Cre systems such as CD19Cre and CD21Cre mice. The *CD19* promoter specifically directs expression beginning at the earliest stages (B220^low^CD43^mid^) and throughout B lymphocyte development ^27^. CD21Cre begins expression later in development at the immature transitional stage and through mature B cells ^28^. Analysis of WT, *Fnip1^−/−^*, *Fnip1^fl/fl^CD19Cre* and *Fnip1^fl/fl^CD21Cre* mice revealed a stage-specific blocks where *Fnip1* deletion occurs in bone marrow cells (Figure 1A). Flow cytometry analysis using fluorescent conjugated B220 and CD43 antibodies showed that both *Fnip1^fl/fl^CD19Cre* and *Fnip1^fl/fl^CD21Cre* allowed B cells to develop farther to B220^lo^IgM^+^CD43^mid^ immature B stage compared *Fnip1^-/-^*mice. However, *Fnip1^fl/fl^CD19Cre* mice displayed significant decrease in representation of immature B cells, and *Fnip1^fl/fl^CD21Cre* mice were disrupted at the B220^hi^CD43^-^igM^+^long-lived mature recirculating B cell stage (Figure 1B). These results using *Fnip1^-/-^, Fnip1^fl/fl^CD19Cre or Fnip1^fl/fl^CD21Cre* collectively suggest that Fnip1 is required for the development of B cells at multiple stages including the pro-B to pre-B cell, pre-B to immature B, and immature to mature B cell transitions (Figure 1 C; modified from^1^).

**Figure 1.**
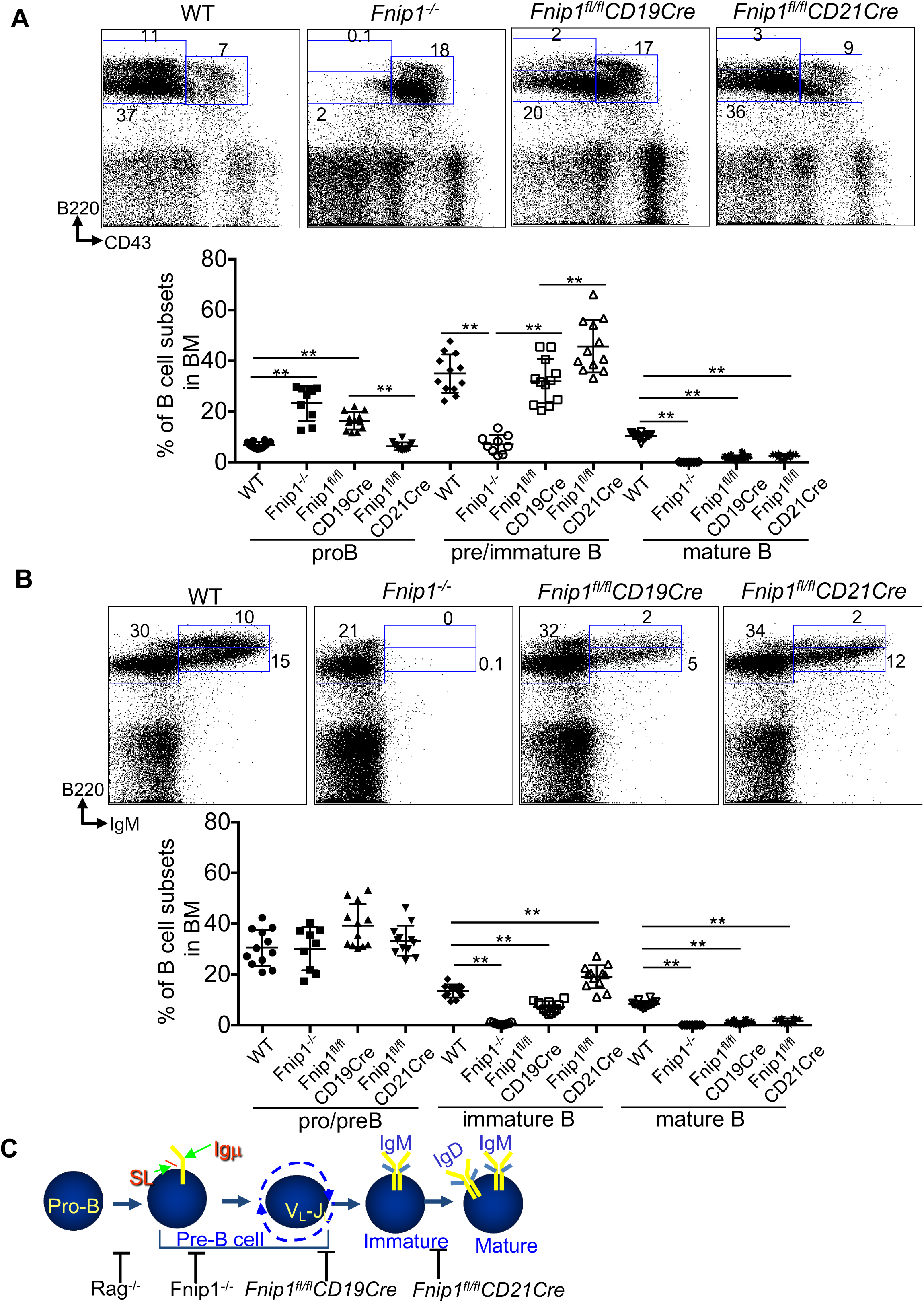
Fnip1 functions as a gatekeeper in B cells development. (A) Stepwise deletion of *Fnip1* during B cell development results in developmental arrest at specific stages, depending on the stage-specific Cre system employed. Flow cytometric analysis of bone marrow (BM) from Fnip1^-/-^ (LPAB.1), *Fnip1^fl/fl^CD19Cre*, *Fnip1^fl/fl^CD21Cre*, and WT mice revealed blockade of B cell progression, as indicated by impaired CD43 downregulation within the B220^+^population. (B) BM cells from *Fnip1*^-/-^, *Fnip1^fl/fl^CD19Cre*, *Fnip1^fl/fl^CD21Cre*, and WT mice were stained with fluorescent-conjugated antibodies and analyzed by flow cytometry. Representative dot plots are shown (n >= 8 mice per group). (C) A schematic (adapted from ^1^) illustrating the stepwise differentiation of B cells, highlighting the differential expression of B220, CD43, and IgM. The stages at which *Fnip1* loss leads to developmental arrest are indicated. Data are presented as mean ± SD; **p<0.01.

### Fnip1 deficiency leads to loss of MZ B cells, reduced FO B cells, and impaired Rag downregulation during B cell maturation

Because *Fnip1^fl/fl^CD21Cre* mice exhibited a loss of recirculating mature B cells in bone marrow, we next examined splenic B cell development. Analysis of splenic cellularity from *Fnip1^fl/fl^CD21Cre* and WT mice showed decreased B220^+^IgM^+^ B cells in the absence of Fnip1 while total numbers of bone marrow and splenocytes were not significantly changed (Figure 2, Figure S1A and S1B). Other immune compartments such as lymph node (LN) and peritoneal cavities contained impaired cellularity of B220^+^IgM^+^ cells in the absence of Fnip1, which suggests the developmental defect by Fnip1 loss leads to impaired B cell maturation (Figure S1C and S1D). By contrast, B1a B lymphocytes that represent a subset of B cells that are found in the peritoneal cavities were relatively intact in *Fnip1^fl/fl^CD21Cre* mice (Figure S1D) ^29^.

**Figure 2.**
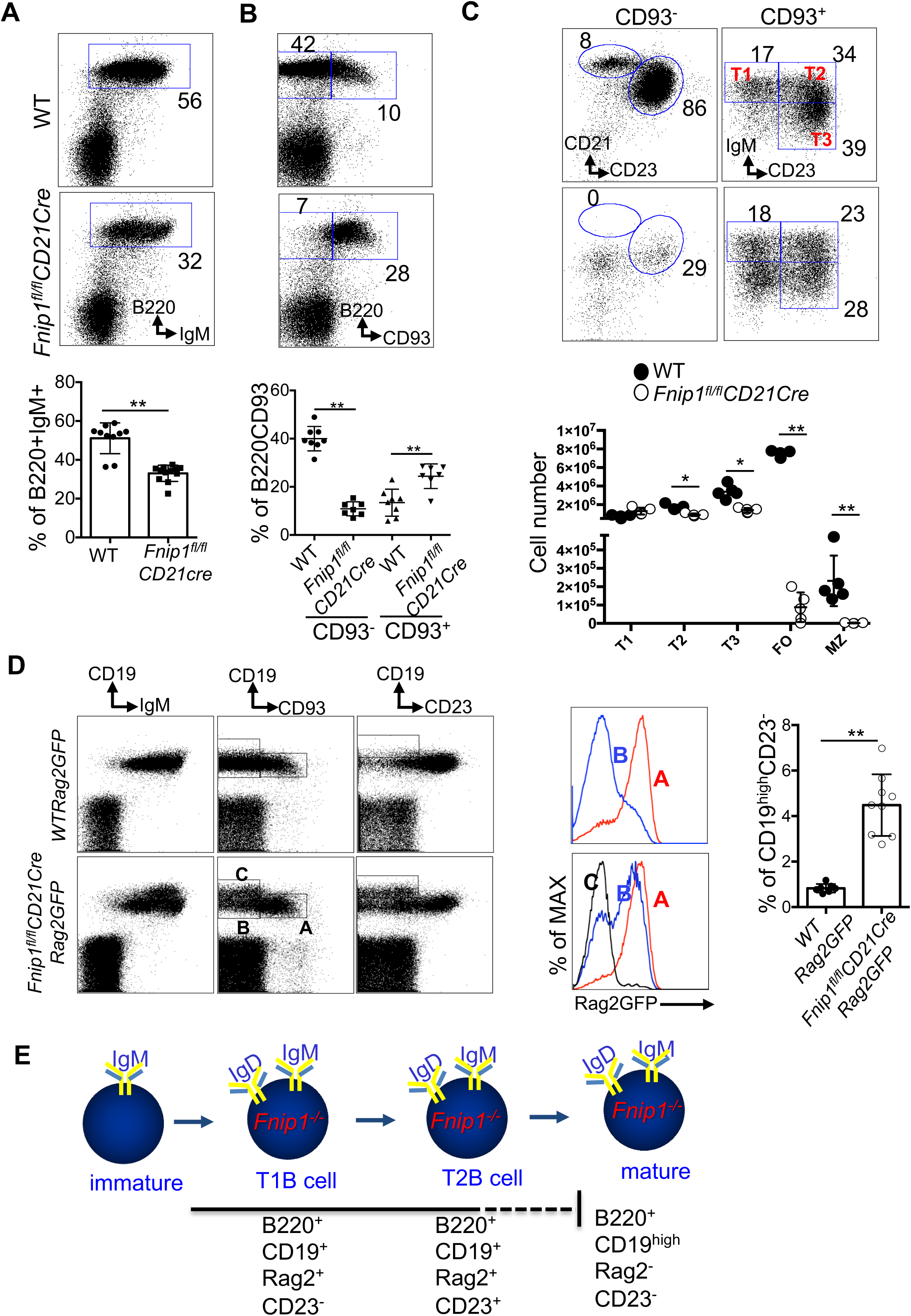
**Fnip1 deficiency arrests splenic B cells at the transitional stage, with persistent Rag activity and high CD19 in *Fnip1^fl/fl^CD21Cre* mice.** (A) Splenic B cell cellularity was markedly reduced in *Fnip1^fl/fl^CD21Cre* mice. Representative flow cytometry plots and the frequencies of B220^+^IgM^+^ cells are shown. (B) *Fnip1^fl/fl^CD21Cre* mice exhibited a developmental block at the transitional B cell stage, as evidenced by increased frequency of B220^+^CD93^+^cells. (C) Maturation of splenic B cells into marginal zone (MZ) and follicular (FO) subsets was impaired in *Fnip1^fl/fl^CD21Cre* mice, which were arrested at the transitional stage. Splenic B cells were stained with fluorescent-conjugated antibodies against B220, CD93, IgM, CD23 and CD21, and analyzed by flow cytometry. Representative plots and absolute numbers of B220^+^IgM^+^cells are shown (n = 5 per group). (D) Splenic B cells from *WTRag2GFP* and *Fnip1^fl/fl^CD21CreRag2GFP* mice were stained with fluorescent-conjugated antibodies against CD19, CD93, IgM, and CD23, and analyzed by flow cytometry (n = 9). Representative flow cytometry plots are shown (left). Rag2 expression in stage-specific splenic B cell subsets is presented (middle), and the frequencies of CD19^hi^CD23^-^ cells are quantified (right). (E) The schematic illustrates the expression patterns of CD19, CD23, and Rag2 during B cell development in Fnip1-deficient splenic B cells. *Fnip1*-deficient splenic B cells exhibit a distinct CD19^hi^Rag2^-^CD23^-^ phenotype prior to their transition into mature B cells. Data are presented as mean ± SD;*p<0.05, **p<0.01.

The reduction in splenic B cell populations observed in the absence of Fnip1 suggests that this phenotype arises from a developmental and/or survival defect during the transitional stage of B cell maturation (Figure 2A). Transitional and mature B cells are typically identified by CD93 expression within the B220⁺ population ^6^. Analysis of CD93 expression revealed that splenic B cells from *Fnip1*^fl/fl^*CD21*Cre mice predominantly exhibit a B220⁺CD93^mid^ phenotype, in stark contrast to wild-type (WT) controls (Figure 2B). Further analysis using fluorescent-conjugated antibodies against IgM and CD23 within the B220⁺CD93⁺ population showed that transitional B cell subsets are differentially represented in *Fnip1^fl/fl^CD21Cre* mice compared to WT mice. In particular, unique subset of B220^+^CD93^+^IgM^low^CD23^-^ (“preTrans B cells”) was enriched in spleen from *Fnip1^fl/fl^CD21Cre* mice (Figure 2C).

To determine whether pre-transitional (T0) B cells originate from the bone marrow in *Fnip1^fl/fl^CD21Cre* mice where B cell development is blocked, we analyzed transitional B cells in the bone marrow, blood, and spleen of both *Fnip1^fl/fl^CD21Cre* and WT mice (Figure S2A). As expected, T0 B cells were present in the bone marrow of both genotypes. However, upon exiting the bone marrow, T0 B cells were no longer detected in the blood or spleen of WT mice, while they persisted in *Fnip1^fl/fl^CD21Cre* mice. To directly evaluate whether T0 B cells are capable of progressing to later transitional stages, we sorted T0 B cells from *Fnip1^fl/fl^CD21 Cre* mice and transferred them into B cell-deficient recipient mice. As expected, the transferred T0 B cells successfully matured into T1–T3 transitional stages (Figure S2B). To assess the efficiency of CD21Cre-mediated deletion, T0-T3 cells were sorted by flow cytometry and their genomic DNA, along with tail DNA, was subjected to PCR analysis. CD21Cre efficiently deleted the Fnip1 floxed allele in T0 and T1 populations, with reduced deletion observed in T2 and T3 cells, possibly due to some selection against deleted alleles during development (Figure S2C). These results indicate that Fnip1 deletion in immature B cells imposes a significant block at the transition from T0 to T1 B cells. In contrast, the transition to T2/T3 B cells based on CD23 expression is less affected (Figure 2C).

However, marginal zone (MZ) B cells, which develop from the T1 subset through T1-MZP precursors, were entirely absent in *Fnip1^fl/fl^CD21Cre* mice, consistent with a block at the T0 to T1 transition predominantly (Figure 2C). To further determine the presence of follicular (FO) B cells, splenic B cells from *Fnip1^fl/fl^CD21Cre* and WT were stained with antibodies against B220, CD93, CD23 and CD21 markers, showed that FO B cells in these mice are phenotypically abnormal (Figure 2C). Specifically, Fnip1-deficient putative FO B cells expressed B220^+^ CD93^low^ CD23^+^ CD21^low^ and were found with small numbers. This is stark contrast to WT FO B cells expressed B220^+^CD93^-^CD23^+^CD21^high^. Together, these data demonstrate that Fnip1 is critical for proper maturation of splenic B cell subsets, including both MZ and FO B cells.

To assess whether *Fnip1^fl/fl^CD21Cre* mice are capable of mounting antibody responses, *Fnip1^fl/fl^CD21Cre* and WT mice were immunized with T-dependent (KLH) and T-independent (NP-Ficoll) antigens (Figure S3A and S3B). Mice lacking Fnip1 produced significantly lower levels of all serum immunoglobulin isotypes in response to both antigens compared to WT controls. Additionally, after immunization with sheep red blood cells (SRBC), *Fnip1^fl/fl^CD21Cre* mice showed reduced germinal center (GC) formation, although the dark zone to light zone ratio assessed via CD38/CD95 and CXCR4/CD83 staining remained normal (Figure S3C and S3D). However, because CD21Cre is active in both mature B cells and follicular dendritic cells (FDC), results from T-dependent antibody responses cannot exclude contributions from FDC-dependent mechanisms. These data indicate that loss of Fnip1 severely impairs splenic B cell development and cellularity, which leads to impaired antibody production.

To evaluate plasmablast differentiation and IgM secretion, splenic B cells isolated from WT and *Fnip1^fl/fl^CD21Cre* mice were stimulated in vitro with LPS, LPS plus IL-4, or anti-IgM /anti-CD40. Measurement of IgM levels demonstrated that Fnip1 deficiency did not impair IgM production compared with WT controls (Figure S4A). Likewise, although MZ B cells are absent in *Fnip1^fl/fl^CD21Cre* mice, total serum IgM levels are comparable to those of WT mice (Figure S4B). Furthermore, analysis of plasmablast differentiation revealed comparable frequencies of B220^low^CD138^+^ cells in WT and *Fnip1*-deficient cultures following stimulation with LPS or LPS plus IL-4, indicating normal plasmablast generation in the absence of *Fnip1* (Figure S4C). In addition, IgD^+^ B cells sorted from WT and *Fnip1^fl/fl^CD21Cre* mice were cultured with LPS and IL-4, and intracellular IgG1 expression was measured and found to be comparable between two genotypes. These results indicate that *Fnip1*-deficient B cells retain the capacity to differentiate into plasmablasts and undergo class-switch recombination similarly to WT B cells (Figure S4D). However, because the total number of splenic B cells are decreased in vivo (Figure 2C), these results collectively suggest that the total number of plasmablasts are likely correspondingly decreased, resulting in reduced antibody levels.

To assess antigen presentation by Fnip1 deficient B cells to CD4^+^ T cells, B220^+^ B cells purified from WT and *Fnip1^fl/fl^CD21Cre* mice were stimulated with LPS for 2 hours and then co-cultured with CFSE-labeled OT-II CD4+ T cells in the presence or absence of OT-II peptides. CD4^+^ T-cell activation was evaluated after 24 hours by measuring CD25 and CD69 expression (Figure S5A), and T-cell proliferation was assessed at day 4 by CFSE dilution (Figure S5B). Under these in vitro conditions, Fnip1-deficient B cells presented antigen and activated CD4^+^ T cells comparably to WT controls. Although in vivo immunization resulted in reduced antibody production, these findings suggest that the defect is unlikely due to intrinsic impairments in antigen presentation by Fnip1-deficient B cells. Instead, it remains likely that defective B-cell localization and/or incomplete B-cell development in vivo limits productive interactions with CD4^+^ T cells.

Precise regulation of V(D)J recombination activating gene (Rag) expression is essential for proper B cell development ^30^. Rag activity is required not only during early bone marrow stages for productive Igμ and Igκ/λ rearrangement, but also in immature B cells undergoing receptor editing of Igκ/λ light chains during transitional B cell development ^31^. Previous studies using a *Rag2*-GFP reporter system have shown that transitional B cells (B220⁺CD93⁺) are uniformly GFP⁺, while mature B cells (B220⁺ CD93⁻) no longer express GFP, indicating successful termination of Rag activity ^32^. To test whether the developmental arrest of transitional B cells in *Fnip1^fl/fl^CD21Cre* mice is associated with persistent Rag expression, we crossed these mice with *Rag2*GFP reporter mice to generate *Fnip1^fl/fl^CD21CreRag2GFP* animals. Consistent with previous reports in WT mice, Rag activity was largely confined to transitional B cells (B220⁺CD93⁺) in the spleen, while mature B cells (B220⁺CD93⁻) were GFP⁻, indicating proper downregulation of Rag expression (Figure 2D). In contrast, *Fnip1*-deficient mice showed persistent Rag activity in spleen, with a majority of splenic B cells remaining GFP⁺, even in populations that phenotypically resembled mature B cells. This persistence of Rag expression is consistent with a developmental block at the transitional stage in *Fnip1^fl/fl^CD21Cre* mice. Moreover, analysis based on CD93 expression within the CD19⁺ population revealed that Rag expression was more robust in splenic B cells from *Fnip1*-deficient mice compared to WT. Strikingly, even CD93^low/-^B cells (normally mature) in *Fnip1*-deficient mice retained high Rag expression, whereas WT counterparts had fully extinguished GFP expression. Interestingly, *Fnip1^fl/fl^CD21CreRag2GFP* mice exhibit a unique population of B cells characterized by elevated CD19 expression, which is absent in *WT Rag2GFP* mice. Detailed analysis of Rag2GFP expression confirmed that these CD19^high^B cells in *Fnip1*-deficient mice have successfully terminated Rag expression.

Since WT T0 cells are present at very low frequency in the spleen, *Fnip1^fl/fl^CD21Cre* cells were compared directly with WT T0 cells within the total B-cell compartment (Figure S6A). WT T0 cells were barely detectable, whereas *Fnip1^fl/fl^CD21Cre* T0 cells were readily observed. Further phenotypic analysis of Fnip1-deficient T0 cells revealed two distinct populations (Figure S6B): (a) CD19^high^CD93^mid^ cells and (b) CD19^+^CD93^mid/high^ cells. Notably, CD19^high^CD93^mid^ cells lacked Rag2 expression with excessive cell growth phenotype, whereas CD19^+^CD93^mid/high^ cells were Rag2 GFP positive. Interestingly, the T0 cells expressed an IgM^low^IgD^high^ phenotype, which is classically associated with mature B cells based on IgM and IgD expression patterns (Figure S6A). Therefore, it is tempting to speculate that the CD19^high^CD93^mid^Rag2^-^ population in *Fnip1^fl/fl^CD21Cre* mice may represent putative mature B cells that enter or accumulate within transitional compartments as a consequence of Fnip1 deficiency, potentially during progression toward follicular (FO) or marginal zone (MZ) B-cell fates. Hence given the heterogeneity of the T0 compartment, it is likely that the CD19^+^CD93^mid/high^Rag2^+^ population represents bona fide T0 cells.

Consistent with this interpretation, these cells gave rise to T1, T2, and T3 populations in in vivo transfer experiments, as shown in the Figure S2B.

As illustrated in Figure 2E, the developmental trajectory of mature B cells in *Fnip1^fl/fl^CD21Cre* mice ends with a distinct phenotype of B220⁺CD19^high^Rag2⁻CD23⁻ cells, in contrast to WT B cells, which typically mature into B220⁺CD19^+^Rag2⁻CD23⁺ cells. These findings indicate that loss of *Fnip1* impairs the downregulation of Rag expression, potentially contributing to the failure of transitional B cells to fully mature.

### Splenic B cells proliferate normally but exhibit reduced viability and impaired homeostatic maintenance in the absence of Fnip1 *in vivo* and *in vitro*

We next examined whether Fnip1-deficient splenic B cells display defects in proliferation and survival in response to various B cell stimuli. CFSE-based proliferation assays demonstrated that total splenic B cells from *Fnip1^fl/fl^CD21Cre* mice divide equally well compared to WT B cells in vitro when stimulated with anti-IgM, LPS, or anti-CD40 antibodies (Figure S7A–C). To assess proliferation in vivo, CFSE-labeled splenic B cells from *Fnip1^fl/fl^CD21Cre* (CD45.2) and WT (CD45.1) mice were mixed and transferred into CD45.2 LPAB recipient mice (B cell deficient mice). Upon in vivo stimulation with anti-IgM, *Fnip1^fl/fl^CD21Cre* B cells proliferated normally and comparably to WT cells (Figure S7D). Similarly, BrdU incorporation assays after 24 hours showed equivalent proliferation between WT and *Fnip1*-deficient B cells (Figure 3A, Figure S8). Together with prior data ^21,23^, these findings indicate that B cell developmental defects in *Fnip1*-deficient mice are not due to impaired B cell proliferation.

**Figure 3.**
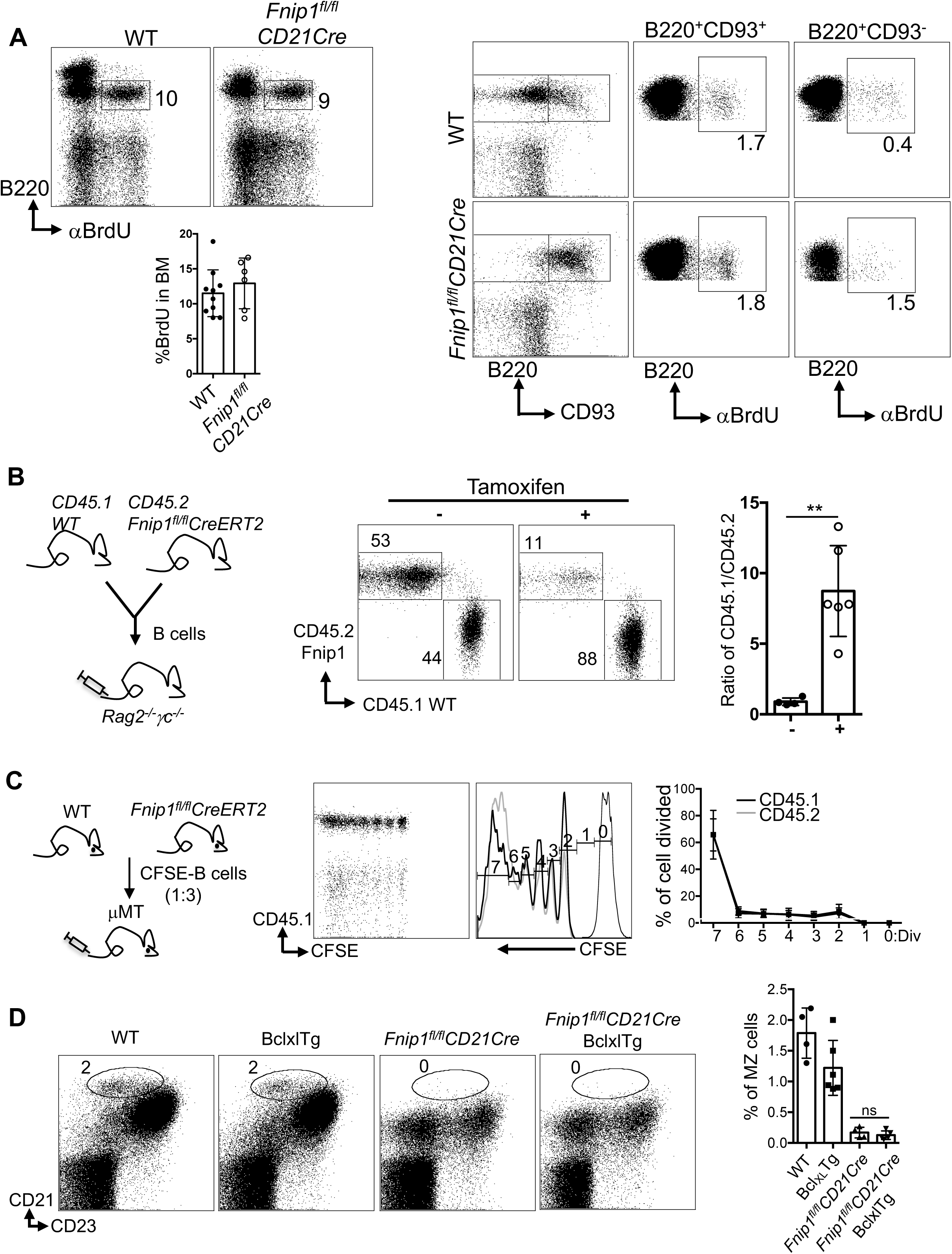
Fnip1 deficiency does not affect B cell proliferation but impairs cell viability and homeostatic maintenance. (A) Proliferation of B cells in the bone marrow and spleen of *Fnip1^fl/fl^CD21Cre* mice was comparable to WT mice. Mice were administered BrdU, and 24 hours later, bone marrow and splenic cells were stained with antibodies against B220, CD93, and intracellular BrdU. Representative flow cytometry plots and quantification of BrdU incorporation are shown, demonstrating normal B cell proliferation in Fnip1-deficient mice. (B) Fnip1 deficiency reduces splenic B cell viability and disrupts homeostatic maintenance. Splenic B cells from *Fnip1^fl/fl^Cre^ERT2^*(CD45.2) and WT (CD45.1) mice were isolated, mixed at a 1:1 ratio, and transferred into Rag2^-/-^γc^-/-^ mice, followed by tamoxifen treatment. The ratio of CD45.1 to CD45.2 cells demonstrates a reduction in the viability and homeostatic maintenance of Fnip1-deficient B cells. (C) Fnip1 deficiency does not affect homeostatic cell division. CFSE-labeled splenic B cells from *Fnip1^fl/fl^Cre^ERT2^* (CD45.2) and WT (CD45.1) mice were mixed at a 3:1 ratio and transferred into μMT mice, followed by tamoxifen treatment. Cell division was assessed by CFSE dilution, demonstrating that Fnip1 deficiency does not impact homeostatic B cell division. (D) Provision of anti-apoptotic signals does not rescue the developmental block. Forced expression of Bcl_XL_ in *Fnip1^fl/fl^CD21Cre* mice did not restore B cell development. The arrested developmental phenotype associated with Fnip1 deficiency was still observed, indicating that the defect is not solely attributable to impaired apoptosis regulation. Data are presented as mean ± SD; **p<0.01.

However, previous work indicate that Fnip1-deficient pre-B cells exhibit significantly reduced viability compared to WT counterparts. To test whether Fnip1-deficient splenic B cells have reduced cell viability, we cultured transitional B cells (CD45.1 WT:CD45.2 *Fnip1^fl/fl^CD21Cre*) enriched with CD93 isolation beads mixed 1:1, and stimulated with or without anti-IgM, anti-CD40, anti-IgM/CD40, and LPS in complete cell culture media. In conditioned media alone, the ratio of CD45.1 WT: CD45.2 *Fnip1^fl/fl^CD21Cre* was significantly increased, indicative of reduced viability of Fnip1 deficient B cells. Anti-IgM alone failed to increase survival of *Fnip1*-deficient cells, whereas anti-CD40, anti-IgM/CD40, and LPS all improved viability, restoring the CD45.1/CD45.2 ratio to levels similar to the input (1:1) (Figure S9A).

We next assessed how nutrient availability affected viability of Fnip1 deficient and WT B cells. We examined their survival in culture media lacking glucose, glutamine, or amino acids. Transitional B cells were first enriched from *Fnip1^fl/fl^CD21Cre* (CD45.2) and WT (CD45.1) mice. A 1:1 mixture of transitional B cells was then cultured in nutrient-rich media or nutrient-deficient media upon anti-CD40 stimulation. After three days, flow cytometry revealed that CD45.2 Fnip1-deficient transitional B cells exhibited increased loss of cellularity compared to CD45.1 WT cells in the absence of amino acids whereas they maintain cellularity comparable to WT in the absence of glucose, and glutamine based on CD45.1/CD45.2 ratios (Figure S9B). These results suggest that Fnip1 is required to maintain metabolic fitness of transitional B cells in response to amino acid availability.

Given Fnip1 role in cell survival and the marked loss of mature splenic B cells in *Fnip1^fl/fl^CD21Cre* mice, we hypothesized that impaired survival also contributes to defective mature B cell homeostasis. To test this notion, splenic B cells were isolated from *Fnip1^fl/fl^CreErt2* (CD45.2) and WT (CD45.1) mice and 1:1 mixed cells were transferred into *Rag2^−/−^γc^−/−^* recipient mice, followed by tamoxifen treatment. The ERT2 system allows Cre expression to be induced by tamoxifen^33^. Ten days later, analysis of flow cytometry showed a significant loss of CD45.2 B cells, while vehicle-treated mice (control) maintained equal ratios of CD45.2 and CD45.1 B cells (Figure 3B). These results collectively suggest that loss of Fnip1 compromises B cell survival.

To further examine homeostatic proliferation, 3:1 CFSE-labeled mixtures of *Fnip1^fl/fl^CreErt2* (CD45.2) and WT (CD45.1) B cells were transferred into μMT mice and treated with tamoxifen. One month later, analysis of CFSE dilution revealed no proliferation defects in Fnip1-deficient cells (Figure 3C), supporting the idea that cell death, rather than proliferation defects, underlies B cell loss. To determine if the loss of viability of Fnip1 deficient B cells is due to Bcl_XL_ modulated survival pathways, *Fnip1^fl/fl^CD21Cre* mice were crossed with *Eμ-Bcl_XL_* transgenic mice, which overexpress the anti-apoptotic protein Bcl_XL_ specifically in B cells (Figure 3D). Bcl_XL_ transgene was unable to rescue development of Fnip1 deficient MZ B cells, suggesting that the developmental and/or survival defect was not dependent on Bcl_XL_ regulated pathways.

To assess apoptosis, B cells from *Fnip1^fl/fl^CD19CreTomato* and *Fnip1^fl/fl^CD21Cre* mice were analyzed by flow cytometry using a Caspase-3/7 FITC assay in combination with a viability (Ghost) dye (Figure S10A and S10B). To enable direct comparison of apoptotic events within the same animals, Tomato^−^ and Tomato^+^ B220^+^CD93^+/-^populations from *Fnip1^fl/fl^CD19CreTomato* mice were stained and analyzed for caspase activity. In parallel, B cells isolated from *Fnip1^fl/fl^CD21Cre* mice were subjected to the same caspase assay. These analyses demonstrated that *Fnip1* deficiency leads to increased apoptosis in the B220^+^CD93^-^ mature B cell population compared with WT controls, whereas *Fnip1*-deficient B220^+^CD93^+^ transitional B cells did not exhibit a significant difference in apoptotic frequency relative to their WT counterparts. Taken together, our findings indicate that Fnip1 is crucial for B cell survival, especially under metabolic stress or BCR stimulation. Loss of Fnip1 disrupts the maturation of B cells, even though their proliferative capacity remains intact.

### Clonal deletion remains intact in Fnip1-deficient B cells

Self-reactive B cells are eliminated or tolerized during early development to prevent autoimmunity ^34^. Immature B cells undergo this selection in the bone marrow and spleen through clonal deletion (apoptosis) and receptor editing, which revises B cell specificity via Ig light chain rearrangement. To assess whether Fnip1 deficiency disrupts B cell tolerance, we utilized the HEL (Hen Egg Lysozyme) transgenic system whereby B cells from anti-HEL B cell receptor transgenic mice (MD4) can be transferred into either membrane bound HEL (mHEL) transgenic mice, which render either a strong BCR or moderate BCR signal respectively upon binding Ag ^35^. Thus, we transplanted bone marrow from *Fnip1^fl/fl^CD21CreMD4* or WTMD4 mice into lethally irradiated KLK4 (mHEL) transgenic mice (Figure 4A). In some experiments, we also used mice generated directly through breeding (i.e., *Fnip1^fl/fl^CD21CreMD4mHEL* or *WTMD4mHEL*). Analysis of splenocytes revealed that introduction of the autoantigen mHEL induced a greater reduction in splenic B cell numbers in *Fnip1^fl/fl^CD21CreMD4mHEL* relative to *WTMD4mHEL* mice, suggesting that clonal deletion is highly efficient in response to strong membrane-bound self-antigen (mHEL) (Fig. 4A, rows 1 and 2). Detailed analysis of the bone marrow and spleen revealed that little or no self-reactive immature B cells escaped into the periphery in both genotypes based on anti-B220 and lysozyme staining, indicating effective central deletion.

**Figure 4.**
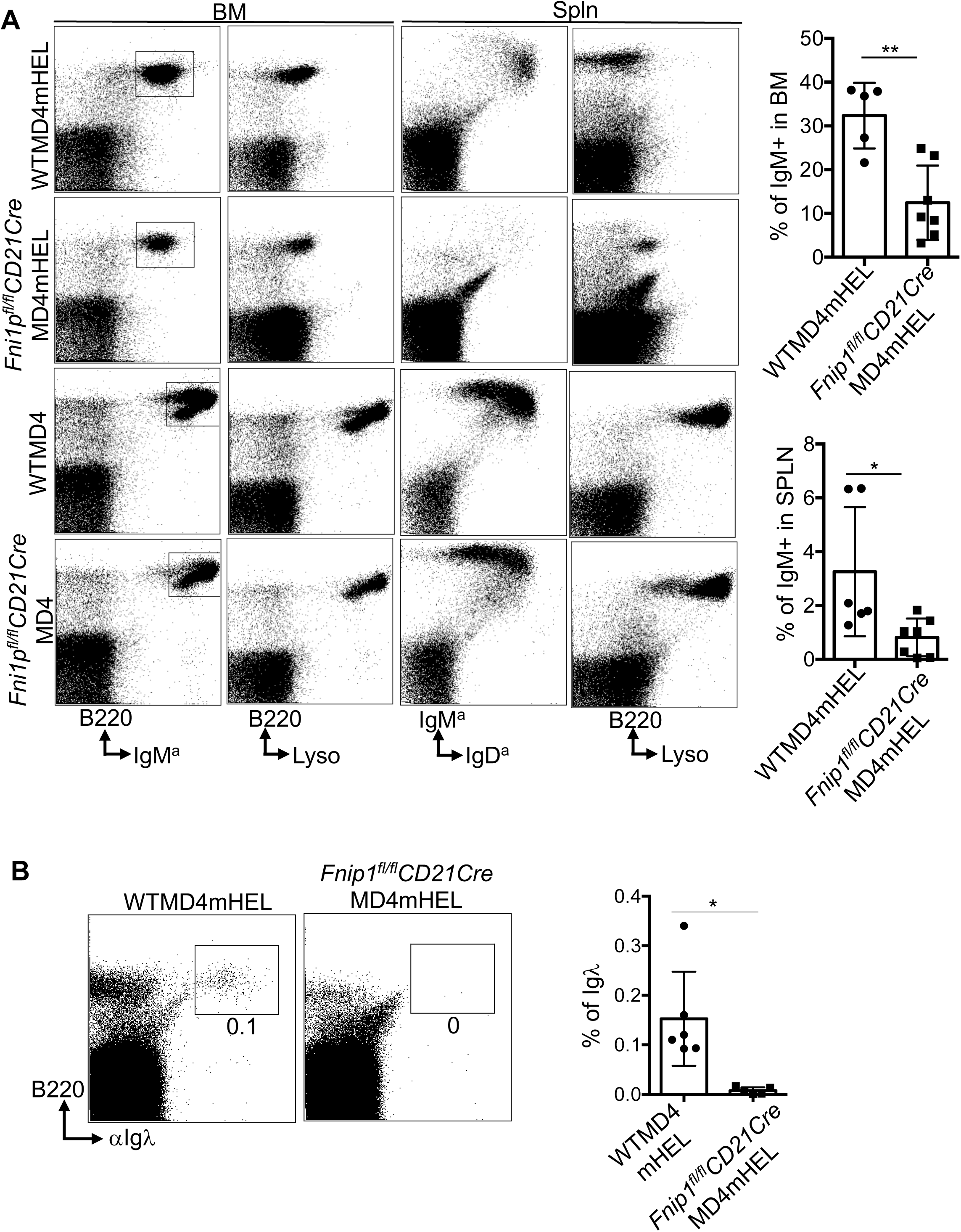
Negative selection is intact in Fnip1-deficient B cells. (A) *Fnip1*-deficient B cells underwent efficient clonal deletion in response to high-affinity antigen (mHEL;KLK4). Bone marrow and splenic cells from *Fnip1^fl/fl^CD21CreMD4mHEL* and *WTMD4mHEL* mice were stained with fluorochrome-conjugated antibodies and analyzed by flow cytometry. Representative dot plots and percentage of cells are shown for bone marrow and spleen from the indicated mice (n=5-7 mice). (B) *Fnip1*-deficient mice exhibited impaired receptor editing and developmental progression compared with WT controls. Splenic B cells were stained with fluorescently conjugated anti B220 or anti-Igλ antibodies and analyzed by flow cytometry. Data are presented as mean ± SD; *p<0.05, **p<0.01.

However, in *WTMD4mHEL* mice, the spleen was partially repopulated with mature IgD^hi^ IgM^lo^ B cells, likely arising through receptor editing, which changes BCR specificity via light chain rearrangement^34^. Consistent with this notion, flow cytometry showed that mature splenic B cells from *WTMD4mHEL* mice did not bind lysozyme (Figure 4A).

In contrast, *Fnip1^fl/fl^CD21CreMD4mHEL* mice lacked these newly edited mature B cells but showed partial accumulation of early transitional B cells (IgM^a+^IgD^a-^). These cells were arrested at the B220⁺CD93⁺IgM^a+^IgD^a-^ stage and may retain low lysozyme reactivity. These results suggest that in the absence of Fnip1, inefficient receptor editing combined with strong negative selection prevents the emergence of non-autoreactive B cells based on anti-B220 and anti-Ig/\ analysis (Figure 4B). Further bone marrow analysis confirmed that immature B cells in both *WTMD4mHEL* and *Fnip1^fl/fl^CD21CreMD4mHEL* mice failed to transition into the IgM^high^ compartment.

Instead, these cells remained IgM^low^ compared to control mice (WTMD4 or *Fnip1^fl/fl^CD21CreMD4*) and retained low lysozyme binding (Fig. 4A, columns 1 and 2), suggesting that BCR signal strength above a critical threshold initiates clonal deletion in this central tolerance model. Interestingly, the level of IgM expression in *Fnip1^fl/fl^CD21CreMD4mHEL* mice, as measured by lysozyme binding, was lower than in WT counterparts, suggesting that self-reactive Fnip1-deficient B cells experience stronger BCR signaling and are more efficiently eliminated (Figure 4A and Figure S11)^36^. Comprehensive analysis of B220^+^Lyso^+^ cells in the bone marrow of *Fnip1^fl/fl^CD21CreMD4mHEL* or *Fnip1^fl/fl^CD19CreMD4mHEL* mice reveals a reduction of these cells in regions where *WTMD4mHEL* mice show a higher presence of B220^+^Lyso^high^ cells (Figure S11A and S11B). This suggests that the BCR signaling threshold is altered in Fnip1-deficient B cells. These data collectively indicate that central tolerance is fully functional in *Fnip1*-deficient B cells and that deletion of autoreactive clones proceeds normally.

### Fnip1 is essential for maintaining peripheral tolerance (anergy)

At later stages of B cell development, peripheral tolerance mechanisms are engaged to prevent activation of self-reactive B cells that escaped central tolerance ^4^. One such mechanism is the induction of anergy, a state of functional unresponsiveness that occurs upon repeated exposure to soluble self-antigens. A hallmark of anergic B cells is the downregulation of membrane-bound IgM (mIgM), reflecting altered BCR surface signaling strength. To investigate whether *Fnip1* deficiency alters the induction of anergy, we transferred bone marrow from *Fnip1^fl/fl^CD21CreMD4* or *WTMD4* mice into lethally irradiated ML5 (sHEL) recipients, that express a soluble form of HEL self-antigen. Analysis of the bone marrow and spleen revealed that WT self-reactive B cells exhibited reduced surface BCR (MD4) expressions based on analysis of IgM and CD23 or B220 and IgM^a^ (Figure 5A and 5B), consistent with an anergic state in *WTMD4sHEL* mice. In contrast, B cells from *Fnip1^fl/fl^CD21CreMD4sHEL* mice maintained higher levels of MD4 BCR expression, comparable to those in *WT MD4*. This indicates a failure to downregulate BCR expression in response to chronic soluble antigen exposure, suggesting impaired anergy induction in the absence of Fnip1.

**Figure 5.**
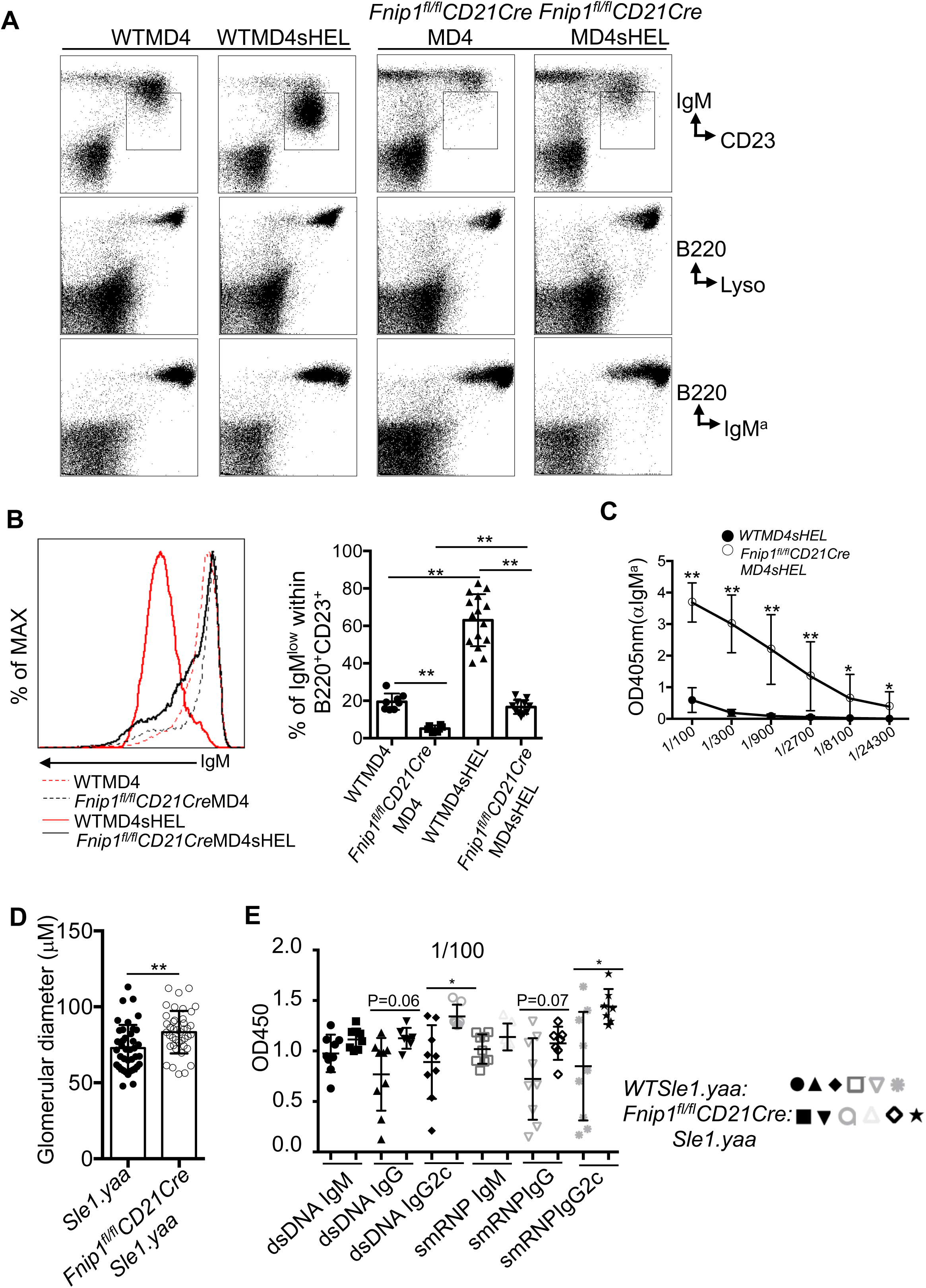
Impaired anergic B cell responses in Fnip1-deficient mice lead to elevated autoantibodies and worsened kidney pathology in an in vivo SLE model. (A) Fnip1-deficient B cells fail to downregulate surface IgM expression in response to low-affinity soluble lysozyme (sHEL; ML5). Bone marrow and splenic cells from *Fnip1^fl/fl^CD21CreMD4sHEL* and *WTMD4sHEL* mice were stained with fluorescent-conjugated antibodies and analyzed by flow cytometry. Representative dot plots with indicated antibody staining are shown. (B) Histogram analysis of IgM expression among *WTMD4*, *WTMD4sHEL*, *Fnip1^fl/fl^CD21CreMD4*, and *Fnip1^fl/fl^CD21CreMD4sHEL* B cells. Bar graph depicts the percentage of B220^+^IgM^low^ cells (boxed in panel A) within the B220^+^CD23^+^ population. (C) Fnip1-deficient B cells produce robust antibody responses against lysozyme. Serum from *Fnip1^fl/fl^CD21CreMD4sHEL*(n=6) and *WTMD4sHEL* mice (n=6) was analyzed for lysozyme-specific IgM^a^ antibodies by ELISA using lysozyme-coated plates. (D and E) Fnip1 deficiency exacerbates kidney pathology and increases autoantibody production in the *Sle1.yaa* mouse model. Glomerular diameter was measured using H&E histology and NDP.view2 software at a 40x magnification (D). Serum autoantibodies were assessed at a 1:100 dilution (E). Data are presented as mean ± SD; *p<0.05, **p<0.01.

To assess functional consequences of defective anergy, we measured lysozyme-specific autoantibodies (IgM^a^ isotype) in serum via ELISA. *Fnip1^fl/fl^CD21CreMD4sHEL* mice produced ∼800-fold higher levels of anti-HEL antibodies compared to *WTMD4sHEL* mice, indicating a breakdown in peripheral tolerance and inappropriate activation of autoreactive B cells (Figure 5C). To test whether Fnip1 loss affects autoimmune disease progression in the *Sle1.yaa* model of systemic lupus erythematosus (SLE), a disease characterized by severe kidney inflammation and elevated autoantibody production due to impaired B cell tolerance^37^, we crossed *Fnip1^fl/fl^CD21Cre* mice with *Sle1.yaa* mice. Histological analysis showed that *Fnip1* loss exacerbated polycystic kidney disease, with enlarged atypical cystic structures and increased glomerulus diameter. This was accompanied by elevated autoantibody levels against dsDNA and smRNP in the serum compared to WT *Sle1.yaa* controls (Figure 5D and 5E and Figure S12). Taken together, these findings suggest that Fnip1 deficiency disrupts immune tolerance and promotes autoimmunity against self-antigens.

### Deletion of CD19 or blockade of the calcineurin/NFAT pathway restores MZ B cells in *Fnip1^fl/fl^CD21Cre* Mice

Because *Fnip1^fl/fl^CD21Cre* mice exhibited elevated expression of the CD19 co-receptor and elevated negative selection indicative of altered BCR signaling, we next sought to determine whether modulation of BCR signaling could restore B cell development in Fnip1 deficient B cells. In particular, CD19 lowers B cell activation thresholds and promotes PI3K/mTOR signaling to regulate B cell development and function ^13,38^. To control BCR specificity and eliminate repertoire differences, *Fnip1^fl/fl^CD21Cre* mice were crossed with MD4 Tg mice. *Fnip1^fl/fl^CD21CreMD4* mice were then bred with *CD19knockinCre* mice in order to delete CD19Cre expression (*Fnip1^fl/fl^CD21CreMD4CD19^d/d^* mice). Flow cytometric analysis revealed that a distinct and significant MZ B cell population in *Fnip1CD21CreMD4CD19^d/d^* emerged one that was absent in *Fnip1CD21CreMD4 and MD4CD19^d/d^* control mice alone. These results suggest that elevated CD19 signaling may be in part linked to impaired MZ development in Fnip1 deficient mice.

We next evaluated the contribution of signaling downstream of CD19, specifically pathways induced by changes in intracellular calcium, and the mTORC1 pathway.

*Fnip1^fl/fl^CD21Cre* mice were treated with cyclosporin A (CSA), a known inhibitor of BCR-induced activation that functions through suppression of Calcineurin activity and nuclear translocation of the transcription factor NFAT, or Rapamycin, a specific inhibitor of mTORC1. Flow cytometry analysis revealed that CSA treatment partially restored the MZ B cell population, suggesting that *Fnip1* may regulate B cell development by modulating BCR signaling strength through the pathways induced by calcium signaling (Figure 6B). In contrast, treatment with rapamycin did not rescue MZ B cell development. This indicates that the impaired maturation of B cells in *Fnip1*-deficient mice is unlikely to result from dysregulated mTOR signaling alone. To further clarify the role of *Fnip1* in the NFAT pathway downstream of BCR signaling, we treated mice with the VIVIT peptide, a specific inhibitor of Calcineurin-mediated NFAT activation. Similar to CSA, VIVIT treatment partially restored MZ B cell development, reinforcing the hypothesis that *Fnip1* influences B cell maturation by modulating BCR signal strength through the Calcineurin/NFAT axis (Figure 6C). Furthermore, to restore MZ B cell development in *Fnip1^fl/fl^CD21Cre* mice, we crossed them with conditional *CnB1^fl/fl^* mice, generating *Fnip1^fl/fl^CD21CreCnB1^fl/fl^* mice. As expected, partial restoration of the MZ B cell population was observed, consistent with the results from CSA and VIVIT treatments (Figure 6D). These findings raise the possibility that *Fnip1* loss shifts the threshold of antigen-dependent BCR signaling.

**Figure 6.**
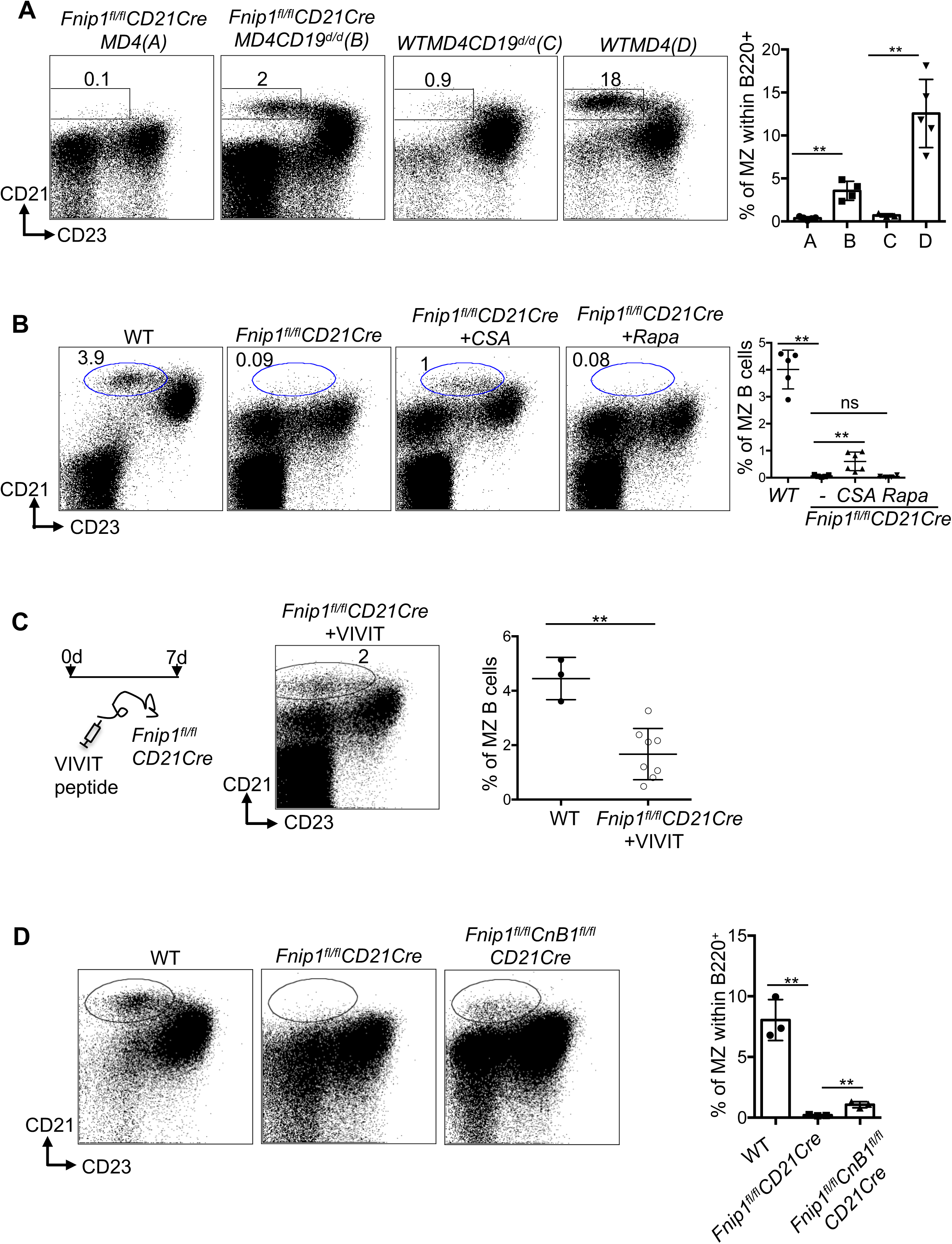
CD19 signaling deletion and calcineurin inhibition partially restore marginal zone B cell development in Fnip1-deficient mice. (A) *Fnip1^fl/fl^CD21CreMD4* mice were crossed with *CD19Cre/Cre (CD19^d/d^)* mice to assess the role of CD19-mediated signaling in marginal zone (MZ) B cell development. Splenic B cells were analyzed by flow cytometry using fluorescent-conjugated antibodies against B220, CD21, and CD23. (B and C) Pharmacological inhibition of calcineurin partially rescues MZ B cells in Fnip1-deficient mice. *Fnip1^fl/fl^CD21Cre* mice were injected intraperitoneally with cyclosporine A (B), rapamycin (B), or a selective NFAT inhibitor (i.v. (C)). Flow cytometry dot plots and quantification of MZ B cells are shown. (D) Conditional deletion of calcineurin in *Fnip1^fl/fl^CnB^fl/fl^CD21Cre* mice partially restores MZ B cell development. Data are presented as mean ± SD; **p<0.01.

### Fnip1-deficient splenic B cells lack nuclear TFEB, show decreased lysosomal content, and display abnormal enlargement

To further elucidate the transcriptional and signaling mechanisms underlying the observed phenotype, cytoplasmic and nuclear fractions were isolated from WT and *Fnip1^fl/fl^CD21Cre* splenic B cells and analyzed by immunoblot. This analysis revealed a marked absence of the transcription factors TFEB and FoxO1 in the nuclear compartment of Fnip1-deficient B cells, indicating impaired transcriptional activation of genes critical for metabolic homeostasis and cell survival (Figure 7A). In parallel, examination of cytoplasmic proteins showed enhanced activation of the PI3K/Akt/mTORC2 and mTORC1 signaling pathways, as evidenced by increased levels of phosphorylated Akt473, phosphor-4EBP and ribosomal protein S6 (pS6R) (Figure 7A). Notably, expression of the metabolic regulator RagD was significantly upregulated in *Fnip1*-deficient B cells, while RagA–C levels remained unchanged, suggesting that the observed defects may be specifically linked to dysregulation of RagD rather than other Rag components ^39^. Consistent with immunoblotting results, real-time PCR analysis revealed that *RagD* expression was markedly increased during the transition from immature to mature B cells in the absence of Fnip1, whereas the expression of other key metabolic regulators and B cell developmental genes remained largely unchanged (Figure S13A). The expression of *RagD* was found to be dependent on BCR stimulation, suggesting that its transcription is regulated by BCR-mediated signaling pathways during B cell development (Figure S13B). Fnip1 is closely linked to the mTORC1/AMPK/FLCN signaling axis, which regulates TFEB nuclear trafficking. To determine whether TFEB-dependent transcriptional programs are altered in the absence of Fnip1, B220^+^CD93^−^ and B220^+^CD93^+^ B cell populations were sorted and analyzed by quantitative real-time PCR. Comparative analysis via qPCR between WT and Fnip1-deficient B cells revealed either normal or slight increases basally in the expression of genes associated with mitochondrial function (atp6v1a, atp6v0a1, nrf1,PGC1alpga), lysosomal biogenesis (mcoln1, ctsd), or autophagy (becn1, sqstm1) (Figure S14A-D)). This finding contrasts with the significant upregulation of the *Rragd* gene observed in Fnip1-deficient cells. These results suggest that *Rragd* upregulation is unlikely to be predominantly TFEB-dependent in the context of Fnip1 deficiency and may instead occur through an AMPK/FLCN-independent mechanism.

**Figure 7.**
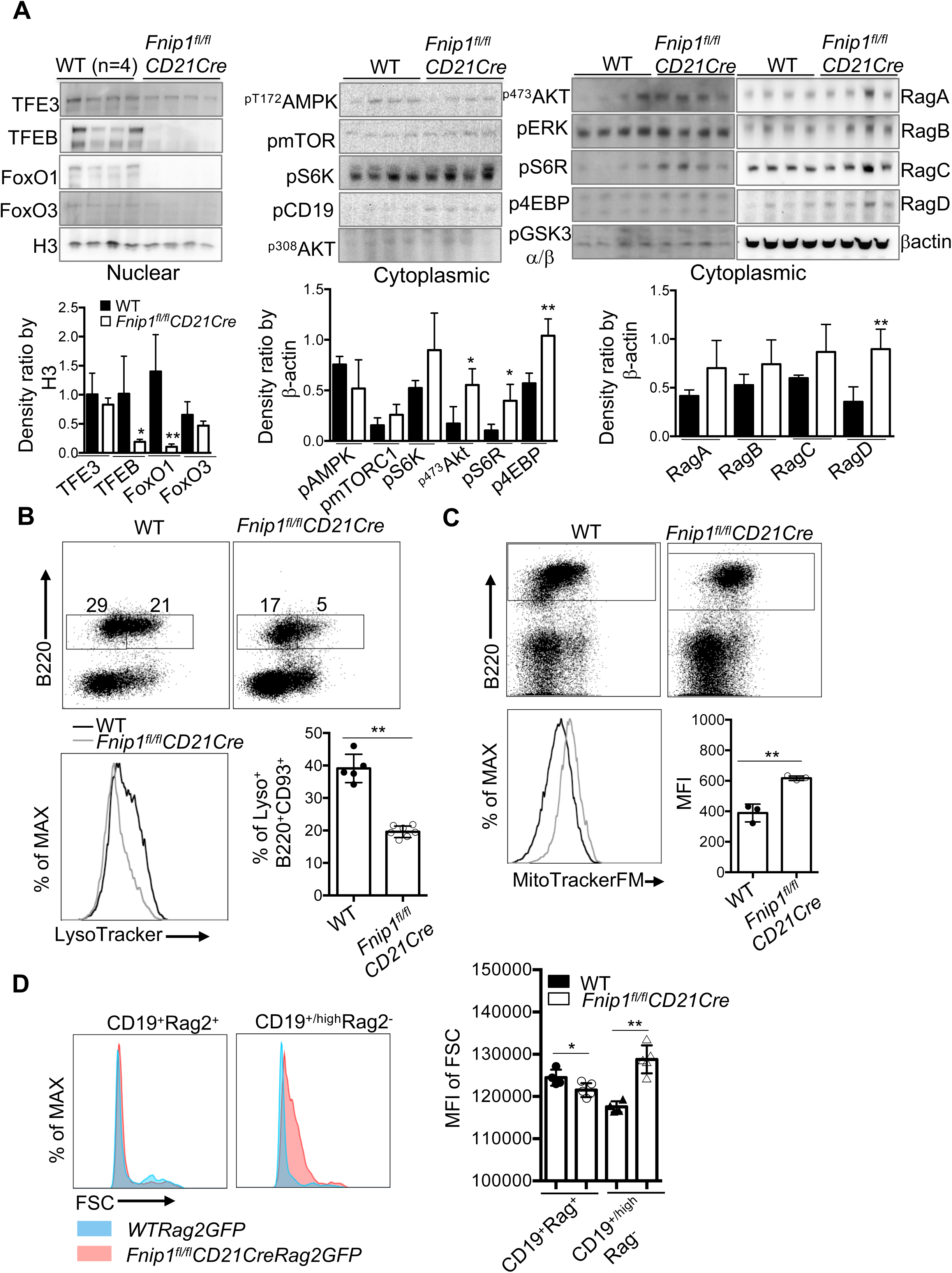
Fnip1 deficiency enhances AKT/mTOR signaling and disrupts TFEB nuclear localization in splenic B cells. (A) Immunoblot analysis was conducted on cytoplasmic and nuclear fractions isolated from B220^+^CD93^+^ B cells purified from WT and *Fnip1fl/flCD21Cre* mice. Enhanced phosphorylation of AKT (pS473), and downstream mTOR targets (pS6R, p4EBP) was observed in Fnip1-deficient B cells, whereas TFEB nuclear localization was impaired. Histone H3 and β-actin were used as loading controls for nuclear and cytoplasmic fractions, respectively. (B and C) Lysosomal and mitochondrial contents in splenic B cells were assessed by flow cytometry using LysoTracker Red DND-99 (B) and MitoTracker FM (C), respectively. Cells were co-stained with fluorochrome-conjugated antibodies against B220. Representative dot plots and histograms are shown, and quantification of LysoTracker and MitoTracker is presented in bar graphs, demonstrating altered organelle content in Fnip1-deficient B cells. (D) B cells undergoing BCR rearrangement in Fnip1-deficient mice exhibit significantly larger cell size compared to their WT counterparts. Cell size was determined by forward scatter analysis. Data are presented as mean ± SD; *p<0.05, **p<0.01.

To validate these molecular findings at the organelle level, we performed flow cytometric analyses using LysoTracker and MitoTracker dyes to quantify lysosomal and mitochondrial content, respectively, in peripheral B cells. Fnip1-deficient B cells exhibited a significant reduction in lysosomal mass, as indicated by diminished LysoTracker staining, and a concurrent increase in mitochondrial content, reflected by elevated MitoTracker signal, relative to wild-type controls (Figure 7B and 7C). Analysis of forward scatter (FSC), which reflects cell size, revealed that Fnip1-deficient CD19^high^Rag⁻ B cells (putative mature B) abnormally become a larger size, whereas their wild-type counterparts exhibited a reduction in cell size during the transition from immature CD19⁺Rag⁺ to mature CD19⁺Rag⁻ B cells (Figure 7D). These findings suggest an imbalance in organelle biogenesis or turnover, consistent with altered metabolic regulation.

## DISCUSSION

Immature B cells undergo a selection process that neutralizes or eliminates self-reactive clones, while those with low or no self-reactivity migrate to the spleen where they continue their maturation into follicular (FO) or marginal zone (MZ) B cells ^6,31^. This progression is tightly coupled with B cell receptor (BCR) signaling, including both tonic and antigen-induced signals, and is critically dependent on the cell’s metabolic fitness to support the energetic demands at each developmental stage ^40^. In this study, we show that loss of Fnip1 expression in transitional B cells results in developmental arrest at the transitional T0/T1 to T2 stages of development. This arrest is due to the inability of Fnip1 deficient B cells to meet the metabolic demands required for proliferation and differentiation, ultimately compromising their viability. In vitro, survival cytokines such as IL-7, SCF, and Flt3L can promote partial progression of Fnip1-deficient pre-B cells to an immature B cell-like state, as evidenced by surface IgM expression ^21^. Consistent with previous findings, while Fnip1-deficient splenic B cells can proliferate, their survival is severely impaired, underscoring the critical role of stage-specific metabolic support for B cell development, even after proliferation ceases.

Fnip1-deficient immature B cells exhibit several unique phenotypes. Despite an increase in peripheral deletion of autoreactive B cells in response to strong BCR signaling (mHEL), peripheral tolerance in response to moderate autoreactive BCR signaling (sHEL) is significantly impaired. These cells display lowered BCR signaling threshold through upregulation of CD19, likely as a compensatory mechanism to preserve metabolic fitness in the face of self-reactive antigen. However, this adjustment in BCR signaling threshold may disrupt receptor editing, leading to negative selection even at low IgM expression levels. Furthermore, Fnip1-deficient B cells resist IgM downregulation in response to soluble self-antigens, resulting in elevated signaling and increased autoantibody production. In the *Sle1.yaa* lupus-prone mouse model, Fnip1 deficiency accelerates severe kidney pathology and autoantibody production, reinforcing the critical role of Fnip1 in maintaining immune tolerance. Fnip1-deficient immature B cells accumulate with a pre-transitional phenotype (IgM^low^CD23⁻), a phenotype nearly absent in wild-type counterparts. These cells also exhibit a unique developmental endpoint characterized by the B220⁺CD93^mid^Rag2⁻CD19^high^CD23^-^ phenotype, accompanied by elevated CD19 expression and elevated mTORC activity, which impedes or delays further differentiation into FO or MZ B cells.

Extensive work in HEK293T cells and other studies has shown that Fnip family proteins integrate into the AMPK and mTOR signaling networks, where they interact with Folliculin (FLCN) and Rag GTPases to regulate lysosomal and mitochondrial homeostasis ^18,20,41^. Under metabolic stress, active AMPK phosphorylates Fnip1, resulting in inhibition of its GAP activity towards RagC, leading to the dissociation of RagC, mTORC1, and TFEB from the lysosome. This dissociation allows nuclear translocation of TFEB, a transcription factor that drives lysosomal and mitochondrial biogenesis. TFEB, in turn, stimulates expression of *RragC* and *RragD*, resulting in feeback non-canonical activation of mTORC.

TFEB is a central BCR controlled rheostats that marks antigen experienced B cells, and drives activation-induced apoptosis in the absence of costimulation^42^. In naïve B cells, TFEB is predominantly localized in the cytoplasm, whereas BCR engagement induces its nuclear translocation. We show that loss of Fnip1 results in increased apoptosis in CD93^-^ B cells while CD93^+^ B cells are largely unaffected (Figure S10). CD40 costimulation partially rescues BCR-induced cell death in Fnip1-deficient B cells (Figure S9), suggesting that cell survival requires additional costimulatory signals to sustain TFEB nuclear localization downstream of defective AMPK/FLCN-dependent TFEB trafficking. In *Fnip1^fl/fl^CD21Cre* mice, CD19^high^CD93^mid^Rag2⁻ follicular and marginal zone B cells may upregulate CD19 and sustain TFEB nuclear localization to compensate. However, elevated non-canonical mTORC1 activity counteracts this response, blocking B cell maturation beyond the T1 stage ^43,44^.

Based on analyses of CD19, CD93, IgM, CD23, and Rag2 GFP expression, we propose that putative mature B cells in Fnip1-deficient mice are arrested at the T0 and T1 transitional compartment. As these cells mature (CD19^high^CD93^mid^Rag2^-^) they acquire enhanced CD19-dependent costimulation, perhaps in attempt to upregulate the AKT/mTOR pathway in order to shut off Rag gene expression via phosphorylation and repression of Foxo1(Figure 7A). However, a detrimental side effect of having elevated CD19 is that it also stimulates increased mTORC1 (which inhibits B cell development at the T1 cell stage), and increased intracellular calcium influx, which activates the phosphatase calcineurin. These results may explain why inhibition of CD19 or calcineurin partially restores B cell development/survival in *Fnip1^fl/fl^CD21Cre* mice (Figure 6).

Interestingly, loss of Fnip1 leads to significant overexpression of *RragD*, while other Rag components remain unchanged or are minimally upregulated, indicating a RagD-specific dysregulation. Elevated RagD may contribute to excessive non-canonical mTORC1 activity in Fnip1-deficient B cells, as has been shown in other tissues such as kidney ^39^. Consistent with this model, elevated RagD expression and increased mTORC1 activity in Fnip1-deficient B cells are associated with enhanced lysosomal sequestration of TFEB^45^, as evidenced by reduced nuclear TFEB levels in B cells from Fnip1^fl/fl^CD21Cre mice (Figure S15). This metabolic profile is further supported by increased mitochondrial content, indicative of a sustained reliance on energetically consuming pathways. Whereas Fnip1-deficient B cells show increased RragD expression and cytoplasmic TFEB, RagA/RagB-deficient B cells exhibit constitutive nuclear TFEB localization and profound humoral immune defects that are rescued by TFEB deletion, implicating Rag GTPases in coupling nutrient sensing to mitochondrial metabolism via TFEB/TFE3 trafficking^41^. These results suggest that Fnip1 deficiency disrupts the balance between AMPK and mTORC1 signaling, favoring anabolic pathways and impairing stress adaptation.

An intriguing feature of *Fnip1*-deficient B cells is the presence of a distinct CD19^high^CD93^mid^RagGFP^-^ population (putative mature B cells) characterized by elevated expression of CD19, a co-receptor that modulates BCR signaling and plays a critical role in B cell development. This upregulation of CD19 is reminiscent of CD19 transgenic mice ^13,38^, which also exhibit breakdown of immune tolerance and increased susceptibility to autoimmunity. In Fnip1-deficient splenic B cells, persistent CD19 expression may alter the BCR signaling threshold, leading to activation of PI3K/mTORC1/mTORC2 and calcium signaling at a lower signaling threshold that disrupts peripheral anergy. Pharmacological inhibition of the Calcineurin/NFAT signaling pathway or genetic deletion of CD19 in Fnip1-deficient B cells partially restores MZ B cell populations, further suggesting that elevated CD19 and calcium signaling axis at least partially contributes to the observed defects in B cell maturation.

Excessive cell growth observed in Fnip1-deficient putative mature B cells (CD19^high^ Rag2^−^) may result from a lowered B cell receptor (BCR) signaling threshold due to elevated CD19 expression. The enhanced activation of the CD19–PI3K–AKT signaling axis promotes continued mTORC1 and mTORC2 activation, resulting in increased cell growth, Foxo1 phosphorylation and nuclear export, and thus *Rag2* silencing. In contrast, wild-type B cells typically exhibit a reduction in mTORC1 and cell size during maturation, transitioning from CD93^+^ to CD93^−^ stages, suggesting that Fnip1 plays a critical role in downregulating mTORC1 signaling during this developmental window.

Consistent with this model, progression to mature B cells requires acquisition of a quiescent metabolic state driven by suppression of mTORC1 signaling. Sustained mTORC1 activation inhibits B-cell development at the T1–T2 transitional stage, as observed in Fnip1-deficient mice ^44^. These findings underscore Fnip1’s importance in restraining aberrant mTORC1 activity and TFEB nuclear location to ensure proper B cell maturation.

The phenotypic and functional abnormalities observed in Fnip1-deficient mice closely mirror the clinical features of human patients with mutations in the *FNIP1* gene, who present with agammaglobulinemia and metabolic dysfunction ^46^. Our findings highlight the importance of Fnip1 in coordinating BCR signaling, metabolic adaptation, and immune tolerance. A better understanding of the molecular mechanisms underlying Fnip1 function and its interactions with key signaling pathways could pave the way for novel therapeutic approaches for immune deficiencies and autoimmune diseases, particularly those linked to metabolic imbalances.

## EXPERIMENTAL PROCEDURES

### Mice

Fnip1floxed, Bcl_XL_ transgenic (*B6.Cg-Tg(LCKprBCL2L1)12Sjk/J*), mHEL (*B6-Tg(KLK4mHEL*)6Ccg, sHEL (*C57BL/6-Tg(ML5sHEL)5Ccg/J*), MD4 (C57BL/6-Tg(*IghelMD4)4Ccg/J*) mice were kindly provided by Dr. Laura Schmidt ^23^, Dr. Pamela Pink, Dr. David Rawlings ^47^ respectively. Additional strains including C57BL/6J (CD45.2), CD45.1 congenic mice (*B6.SJL-Ptprc^a^ Pepc^b^/BoyJ*), *Ighm^tm1Cgn^*, LPAB (Fnip1^-/-^)^21^, *Rag2^−/−^ψc-/-* (*C57BL/6NTac.Cg-Rag2^tm1Fwa^ Il2rg^tm1Wjl^*) mice, Rag2EGFP FVB-Tg (*Rag2-EGFP)1Mnz/J*), CD21Cre (*B6.Cg-Tg(Cr2-Cre)3Cgn/J*), CD19Cre (*B6.129P2(C)-Cd19^tm1(Cre)Cgn^/J*), CnBfloxed (*Ppp3r1^tm2Grc^/J*), Ert2Cre (*B6.Cg-Ndor1^Tg(UBC-Cre/ERT2)1Ejb^/1J*), SLE1yaa (*B6.Cg-Sle1^NZM2410/Aeg^ Yaa/DcrJ*), and *μMT* (*B6.129S2-Ighm^tm1Cgn^/J*) mice were obtained from the Jackson Laboratories or Taconic Biosciences. Fnip1floxed mice were backcrossed to the C57BL/6J background for at least 10 generations. Experimental studies were conducted using mice aged from 8 to16 weeks. Littermate controls were used whenever available. Experimental controls, denoted as WT, include *Fnip1^fl/+^*, *Fnip1^fl+^CD21Cre and Fnip1^fl/fl^* mice. All mice were housed under specific pathogen-free (SPF) conditions in facilities at the University of Washington. All experimental protocols involving animals were reviewed and approved by the Institutional Animal Care and Use Committee (IACUC) of the University of Washington.

### Reagents

A comprehensive list of reagents used in this study is as follows with chemical reagent, dye, immunological reagent, cell culture reagents, antibodies, and magnetic bead based kits: Corn oil (C116025, Aladdin), Bromodeoxyuridine (BrdU, fisher scientific), 4OH tamoxifen (6833585, peprotech), CellTrace™ CFSE Cell Proliferation Kit (C34554, fisher scientific), lysozyme (210083410, MP biomedicals) cyclosporin A (239835, EMD millipore chemicals), rapamycin (R-5000, LC laboratories), NFAT inhibitor (VIVIT, 249537-73-3, cayman chemical), LysoTracker™ Red DND-99 (L7528, thermofisher scientific), MitoTrackerGreen FM (M46750, thermofisher scientific), F(ab’)2-Goat anti-mouse IgM (31178, Invitrogen), αCD40(14-0402-86, fisher scientific), LPS from *Escherichia coli* O111:B4 (L4391, sigma aldrich), MEM Amino Acids Solution (50X) (11130-051, Invitrogen), Indo-1 (I-1223, thermofisher scientific), sheep red blood cells (ISHRBC100,Innovative research), keyhole limpet hemocyanin (374805, EMD millipore chemicals), NP-Ficoll (f1420-10, bioresearch technology), Imject Alum Adjuvant (77161,thermofisher scientific), NP-BSA (n-5050m-10, biosearch technologies), anti mouse IgM HRP (1021-05, southern biotechnology), anti mouse IgG2a HRP (1081-05, southern biotechnology), anti mouse IgG HRP (sc-2005, santa cruz biotechnology), anti mouse IgG1 HRP (1071-05, southern biotechnology), anti mouse IgG3 (1101-05, southern biotechnology), αCD169 (142406, Revvity), BD pharmingen^TM^ FITC brdU kit (559619, BD bioscience), DAPI (H1200, vector lab), B cell isolation kit (130-090-862, miltenyi biotech), CD93 microbeads (130-105-753, miltenyi biotech), BD Cytofix/Cytoperm™ Fixation/Permeabilization Solution Kit (BDB554714, BD Bioscience), RPMI-glucose (11879-020, invitrogen), D-glucose (G8270, sigma aldrich), L-glutamine (21051-024, thermofisher scientific), RPMI-glutamine (21870-076,

Invitrogen), RPMI 1640 Medium GlutaMAX (61870036, thermofisher scientific), fetal bovine serum (97068-085,VWR international), MEM Non-Essential Amino Acids Solution (100X) (11140050, fisher scientific), sodium pyruvate (100 mM)(11360070, thermofisher scientific), MEM vitamin solution 100X (11120-052, Invitrogen), HEPES (1M)(15630106, thermofisher scientific), MEM Amino Acids (50x) solution (M5550-100M, Invitrogen), DPBS (14190-250, Invitrogen).

### Immunoblotting

Immunoblot analysis was conducted as previously described ^21^. Briefly CD93^+^ B cells (5×10^6^) isolated from spleen were lysed and fractionated into nuclear and cytoplasmic compartments using NE-PER™ Nuclear and Cytoplasmic Extraction Reagents (78833, thermofisher scientific) according to manufacturer’s protocol. Immunoblot was probed with primary antibodies against Rabbit polyclonal antibodies or mouse monoclonal antibodies specific for βActin (8457, cell signaling; A5441, sigma Aldrich), P-Ser473 Akt (4060P), P-Thr308 Akt (13038T), P-Ser240/244 S6 ribosomal protein (5364S), P-Thr172AMPK (2535), Phospho-mTOR (Ser2448), Phospho-p70 S6 Kinase (Thr389)(9234S), Phospho-CD19 (Tyr531) (3571T), Phospho-p44/42 MAPK (Erk1/2) (Thr202/Tyr204) (4370P), p44/42 MAPK (Erk1/2) (4695P), FoxO1 (2880S), FoxO3a (2497T), Phospho-4E-BP1 (Thr37/46) (2855S), Phospho-GSK-3α/β (Ser21/9)(9327T), Rag and LAMTOR antibody sampler kit (RagA (4357), RagB (8150), RagC (9480), RagD (4470)), Histone H3 (4499P), TFE3 (14779s), TFEB (A303-673A, bethyl lab), α-rabbit IgG HRP (W4011, promega), α-mouse IgG HRP (W4021, promega), and others were purchased from Cell Signaling. Signals were visualized using chemiluminescent detection (ECL).

### Flow cytometry

Cells were harvested from bone marrow (BM), spleen, lymph nodes (LN), peritoneal cavity, and blood and stained with fluorescent-conjugated antibodies specific for B220 (RA3-6B2, BioLegend), CD43 (S7, BD Pharmigen), IgM (DS-1, BD Pharmigen), IgD (11-26C.2A, BioLegend), CD93 (AA4.1, BioLegend), (AA4.1, BioLegend), CD23 (BEB4, BioLegend), CD21 (7E9, BioLegend), CD19 (1D3, Tonbo Biosciences), CD45.2 (104, Cyteck), CD45.1 (A20, Cyteck), IgMa (MA69, Biolegend), IgDa (AMS9.1, Biolegend) Ig/\ (RML42, Biolegend), biotinylated lysozyme, CD5 (53-7.3, Fisher), CD38 (90, BioLegend), CD95 (Jo2, BD bioscience), CXCR4 (2B11,eBioscience), CD83 (Michel19,Biolegend), fluorophore labeled streptavidin (PE/ APC) were used for antigen specific staining. Data were acquired using flow cytometry as previously described and analyzed with Flowjo software.

### In Vivo homeostatic survival and proliferation assay

Splenic B cells from *Fnip1^fl/fl^CD21Cre* (CD45.2) and WT (CD45.1) were isolated using a negative selection B cell isolation kit. 1:1 mixed cells (1×10^7^) was intravenously into *Rag2^-/-^ψc^-/-^* recipient mice. Tamoxifen (1 mg/mL) was administered via light-protected drinking water to induce gene deletion ^33^. Ten days post-transfer, splenocytes were harvested and the relative proportions of CD45.1 versus CD45.2 cells were determined by flow cytometry. For proliferation assay, CFSE-labeled WT and *Fnip1^fl/fl^CD21Cre* B cells were co-transferred into μMT mice. Four weeks later, proliferation was assessed by CFSE dilution in CD45.1 and CD45.2 B220^+^ cells.

### Pharmacological inhibition studies

Mice were treated intraperitoneally with Cycloporin A (100μg) and Rapamycin (100μg) every other day. VIVIT peptide (20μg) was administrated intravenously twice within a 7-day period. Cyclosporin A was dissolved into pharmaceutical-grade corn oil. Rapamycin in PEG400/Tween 80 vehicle, and VIVIT peptide was dissolved in sterile saline prior to injection.

### BrdU incorporation in Vivo

To assess cellular proliferation, mice were injected intraperitoneally with 100μg BrdU and. 24 hours later, bone marrow and splenic B cells were stained surface markers, fixed, permeabilized, and treated with DNase I (30μg/sample, Sigma-Aldrich). Cells were then stained with FITC-conjugated anti BrdU antibody (BD Biosciences) and analyzed via flow cytometry.

### Enzyme-linked immunosorbent assay (ELISA)

NUNC Maxisorb plates were coated with KLH (50 μg/mL), NP BSA (10μg/ml), or lysozyme (50μg/ml) and incubated overnight at 4°C. After blocking with 10% BSA in PBS, serial dilutions of serum were added. Isotype-specific HRP-conjugated secondary antibodies (IgM, IgG, IgG2a, IgG1, IgG3) were used. Colorimetric detection was performed using Super AquaBlue ELISA Substrate (thermofisher), and absorbance was measured at 405 nm.

### Statistical analysis

All data were analyzed using GraphPad Prism version 6.0. Statistical significance was determined by unpaired two-tailed Student’s *t*-test, as appropriate. A *p*-value less than 0.05 (*) was considered statistically significant.

## Author contributions

HP and BMI designed the experiments and wrote the manuscript. HP, RC, DS, RS conducted the experiments, collected data and performed the analysis.

## Acknowledgements

This work was supported by NIH grants R01AI109020, R01AI158353, and R21AI156243 awarded to BMI. Bridge Funding support was provided by the University of Washington Provost’s Office and the Department of Comparative Medicine.

## Conflict of Interest

The authors declare that they have no competing interests.

---

**Figure S1.**
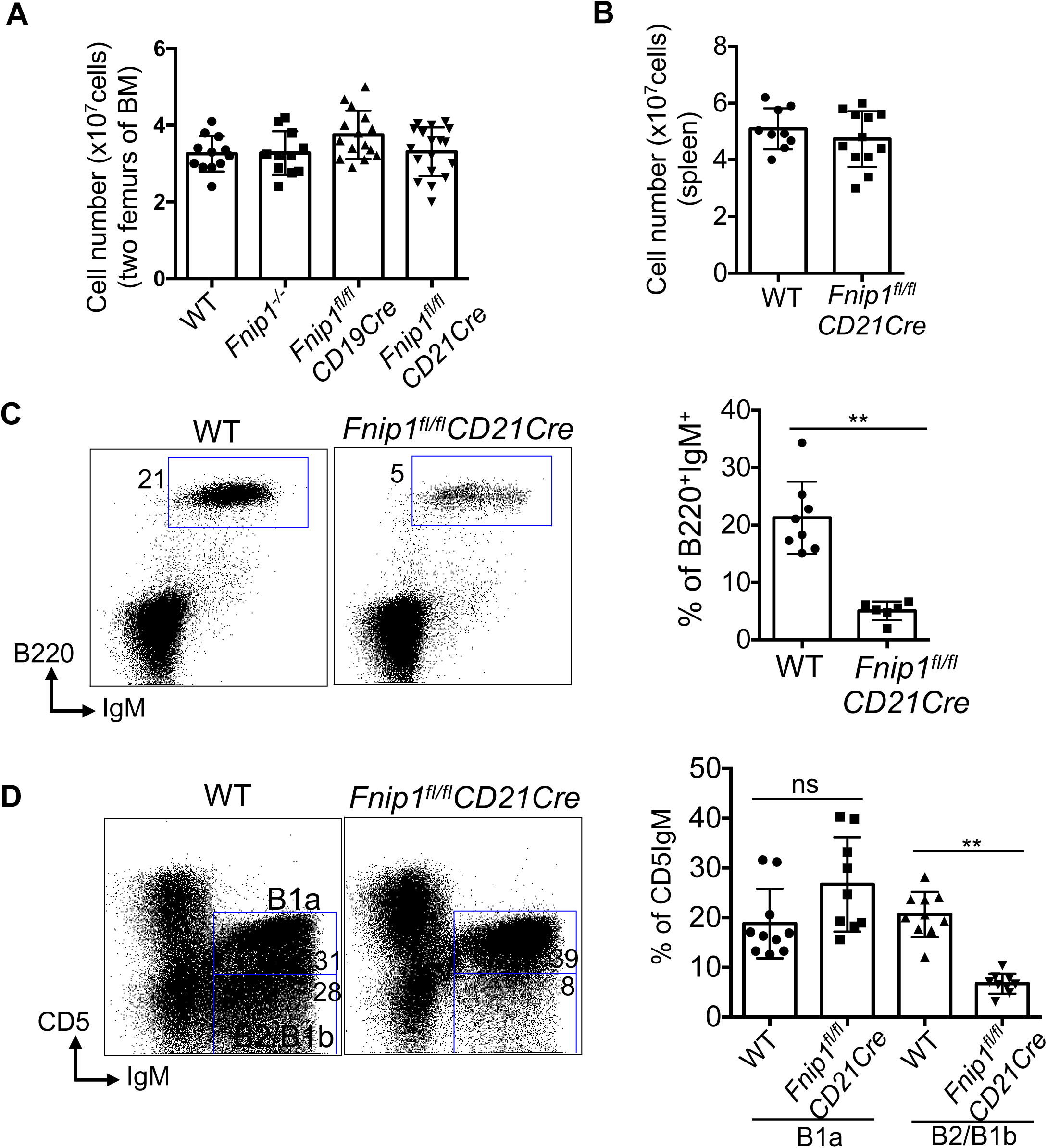

**Figure S2.**
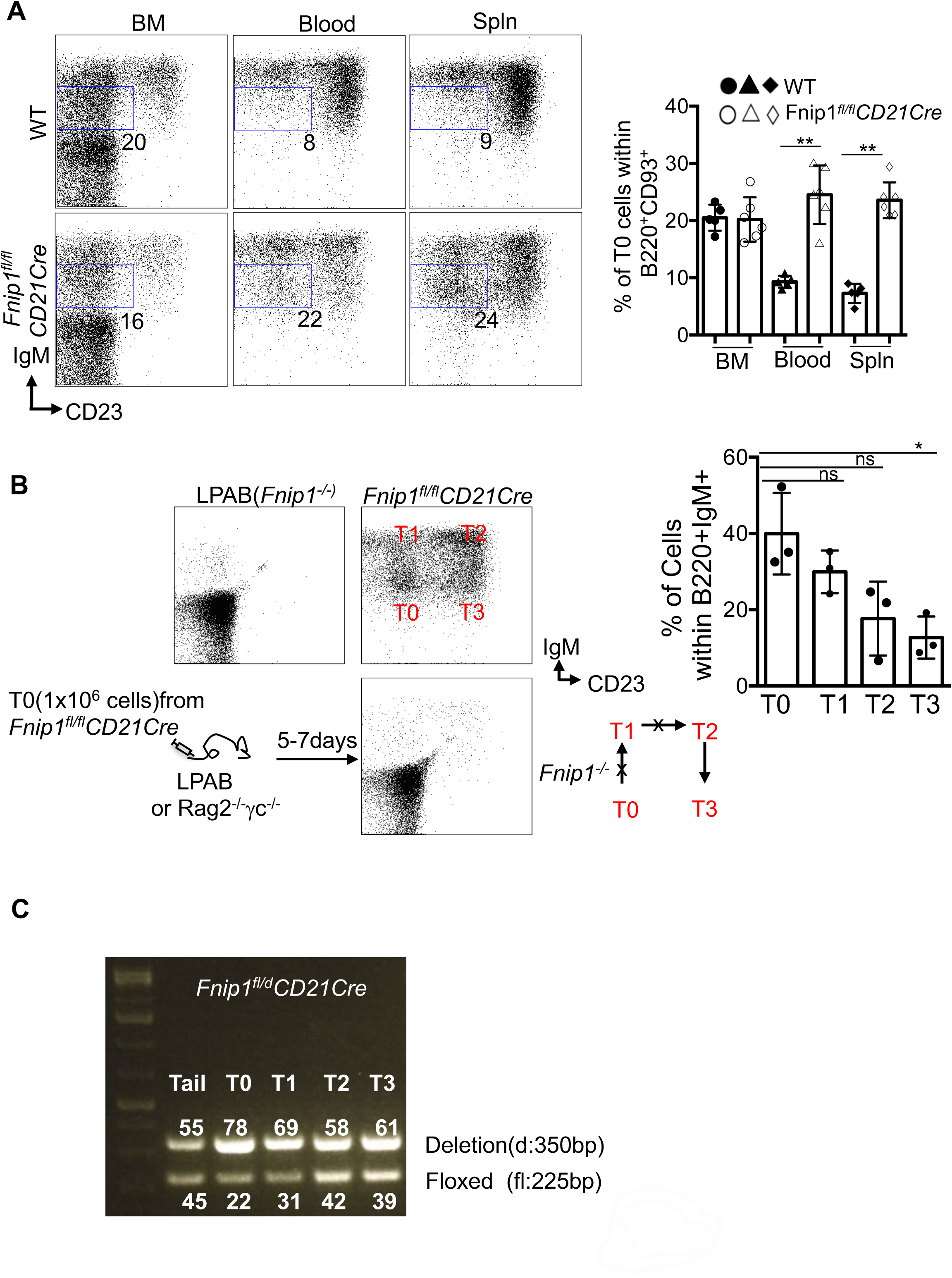

**Figure S3.**
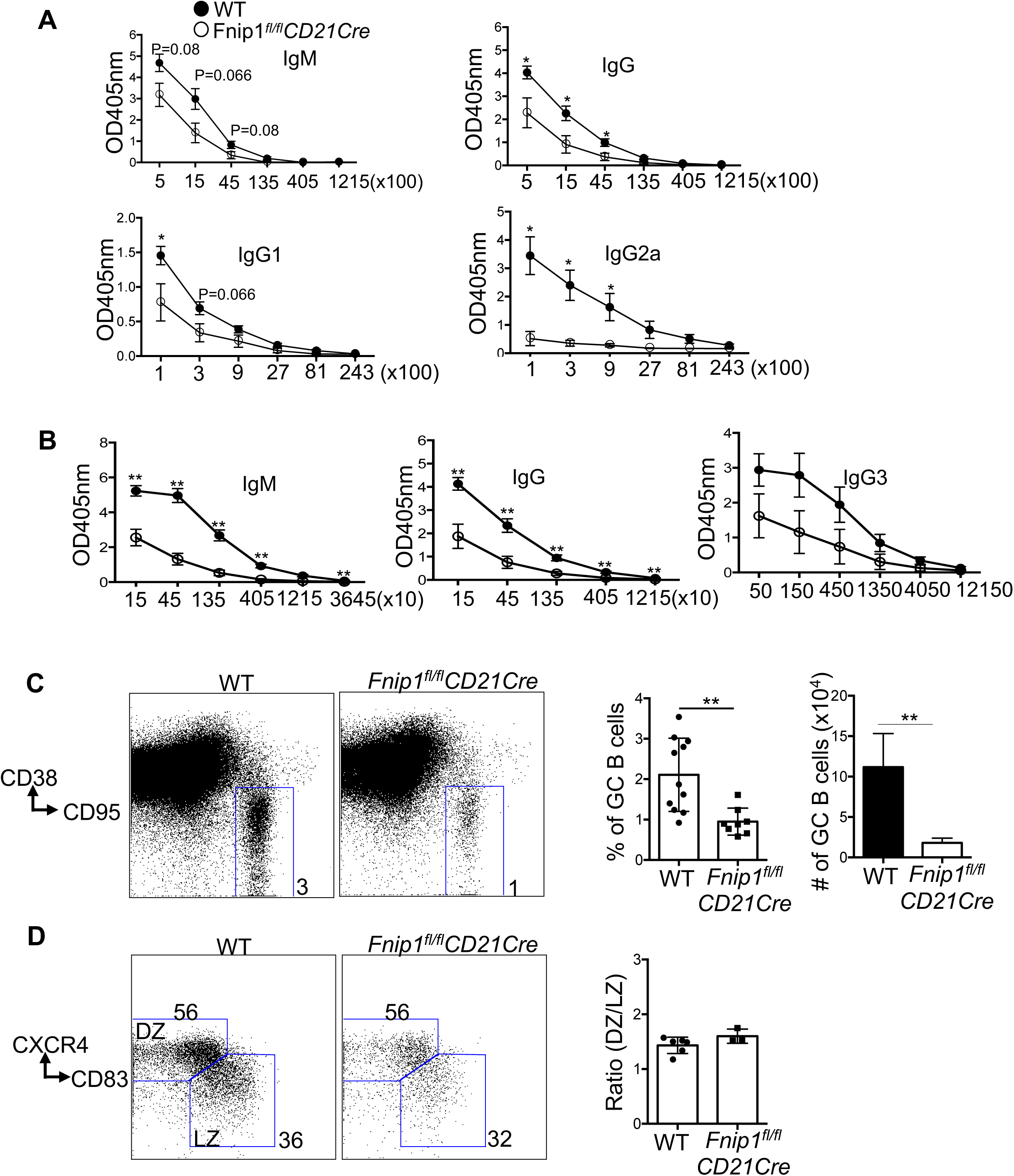

**Figure S4.**
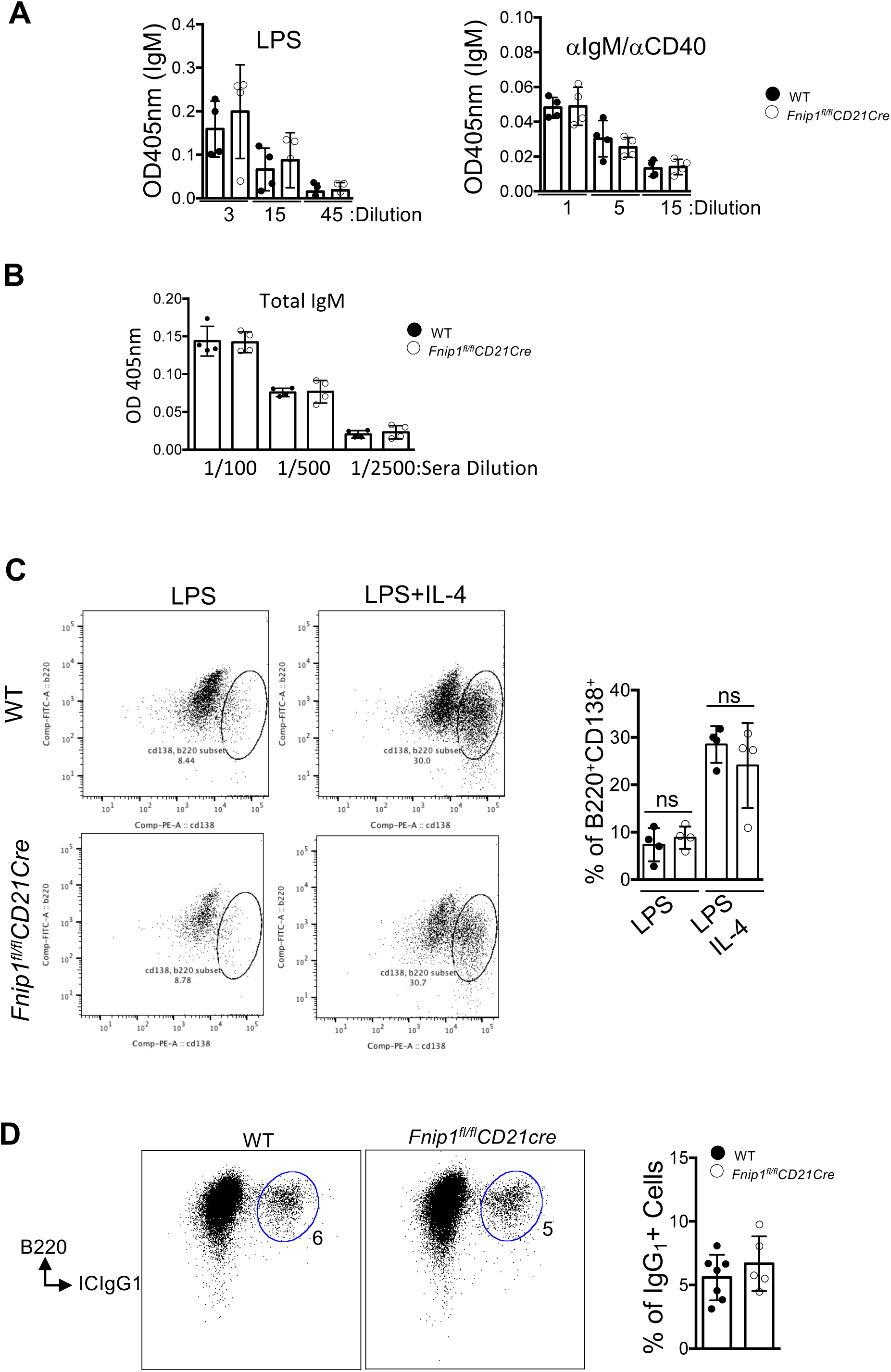

**Figure S5.**
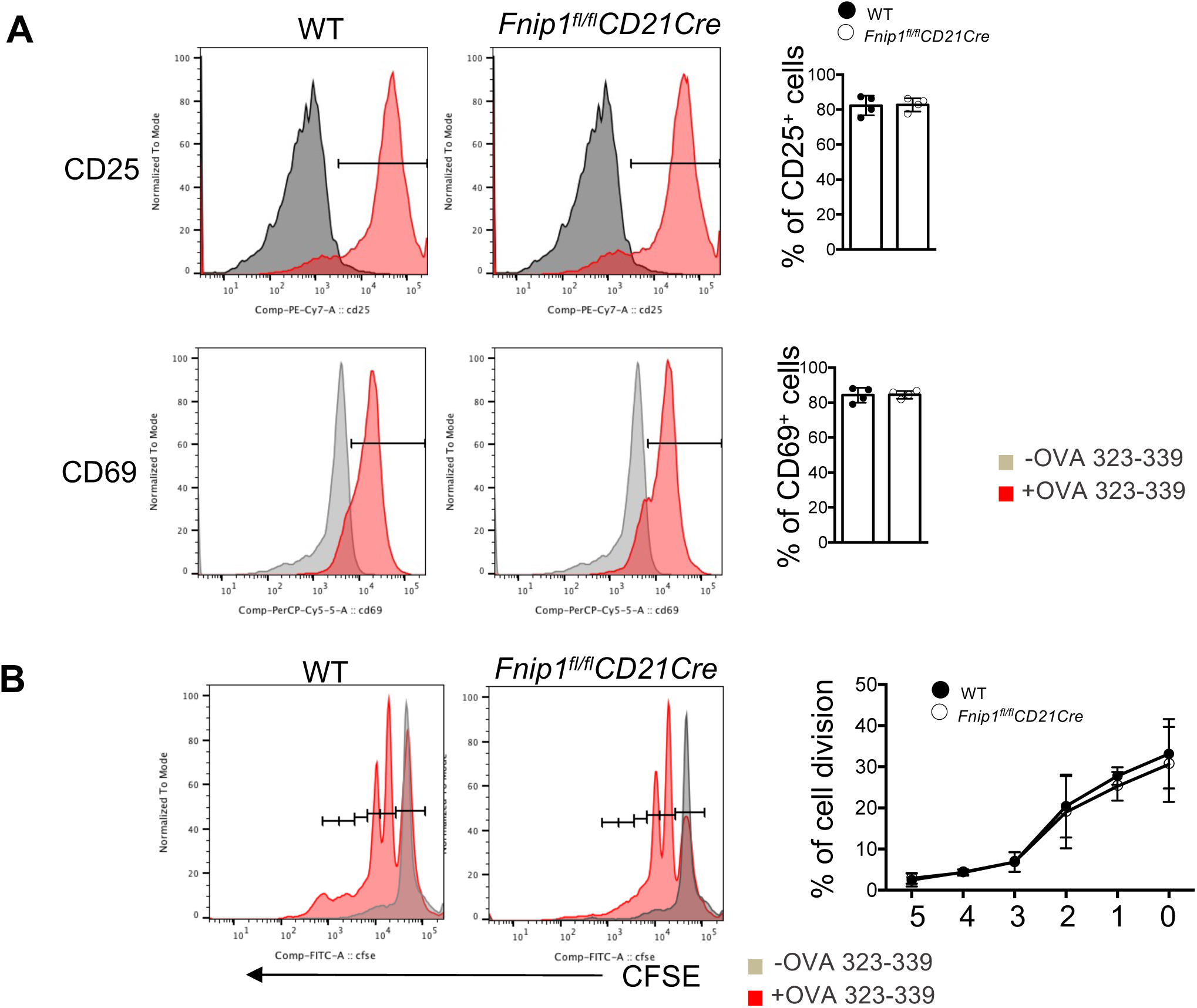

**Figure S6.**
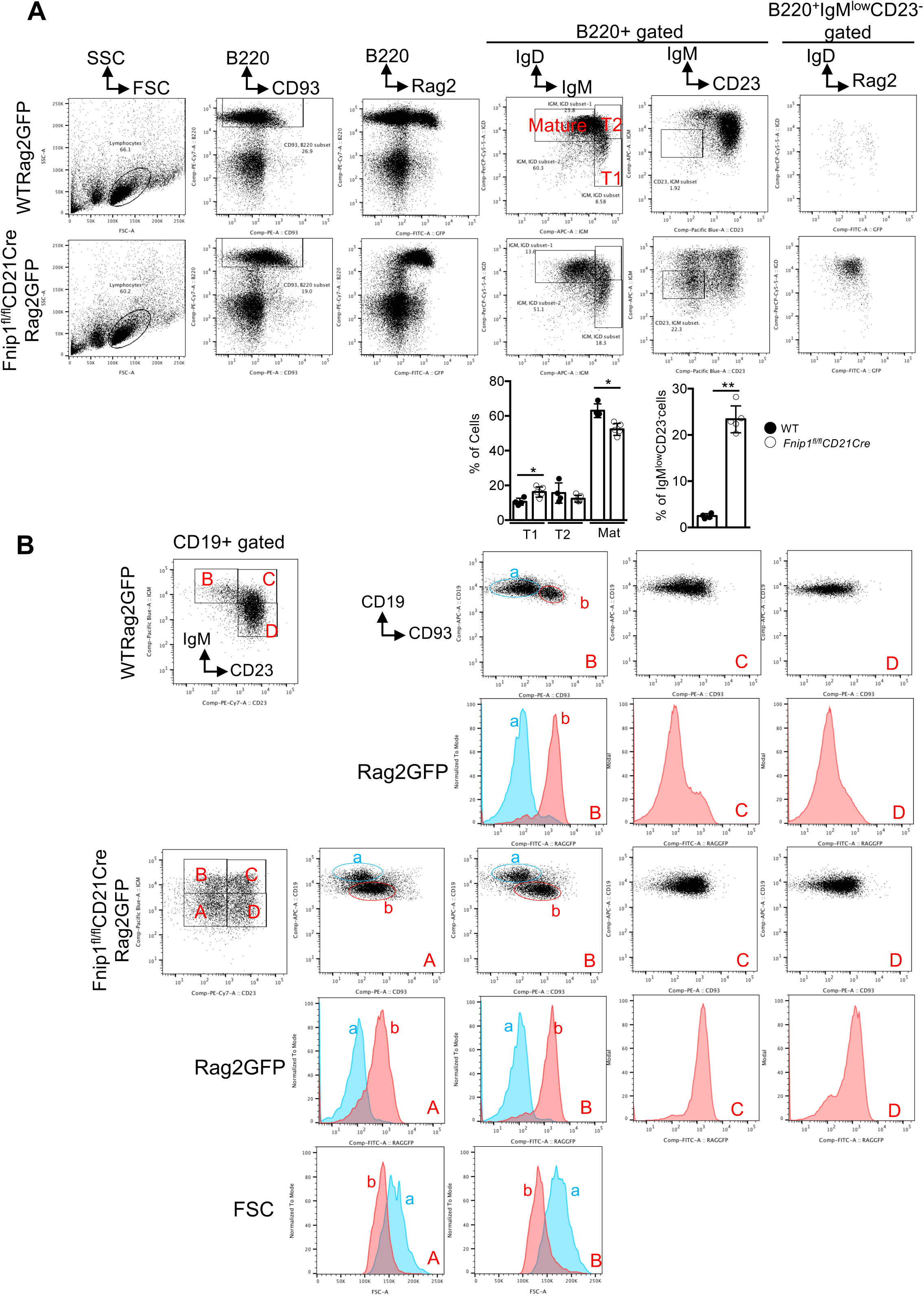

**Figure S7.**
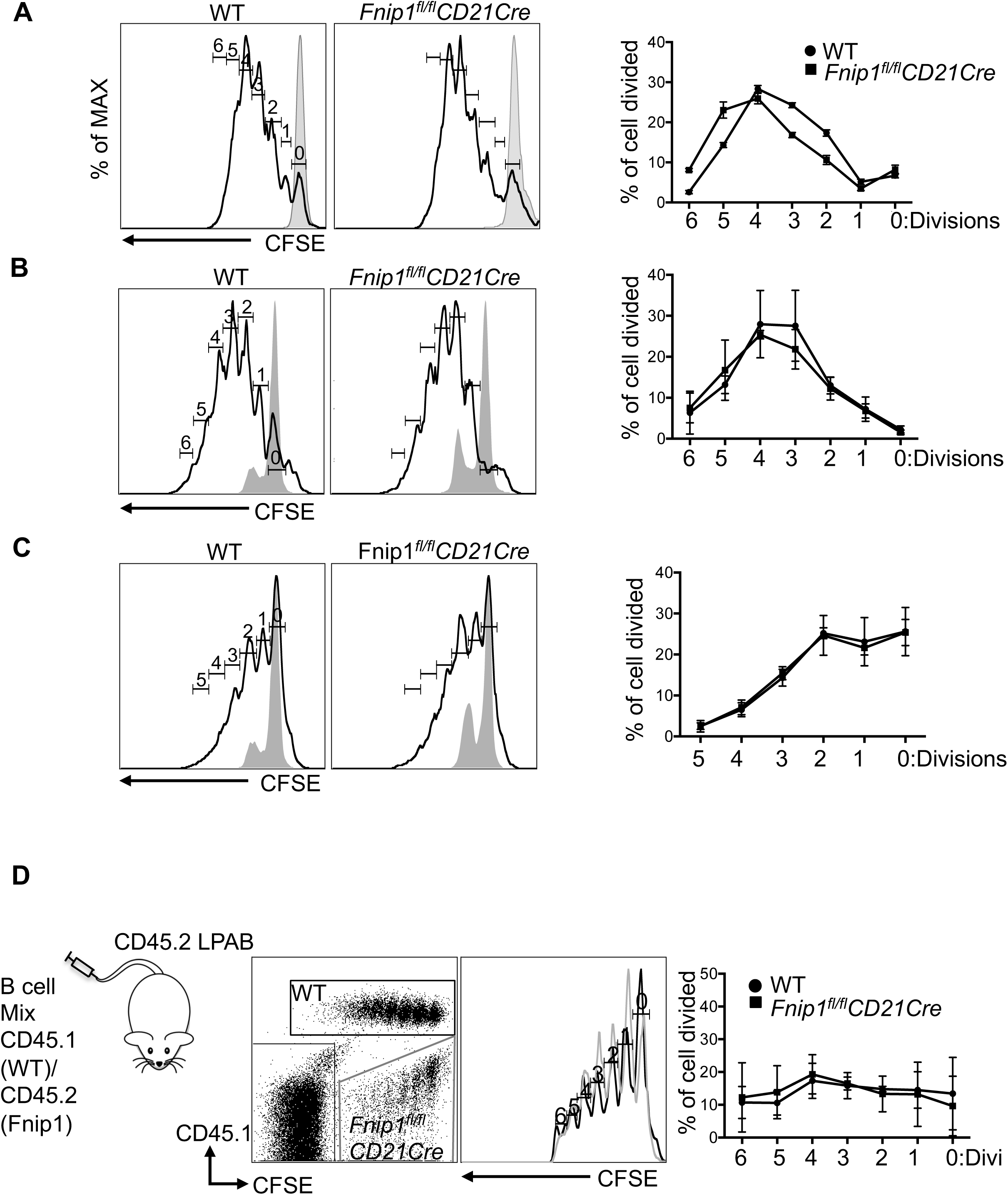

**Figure S8.**
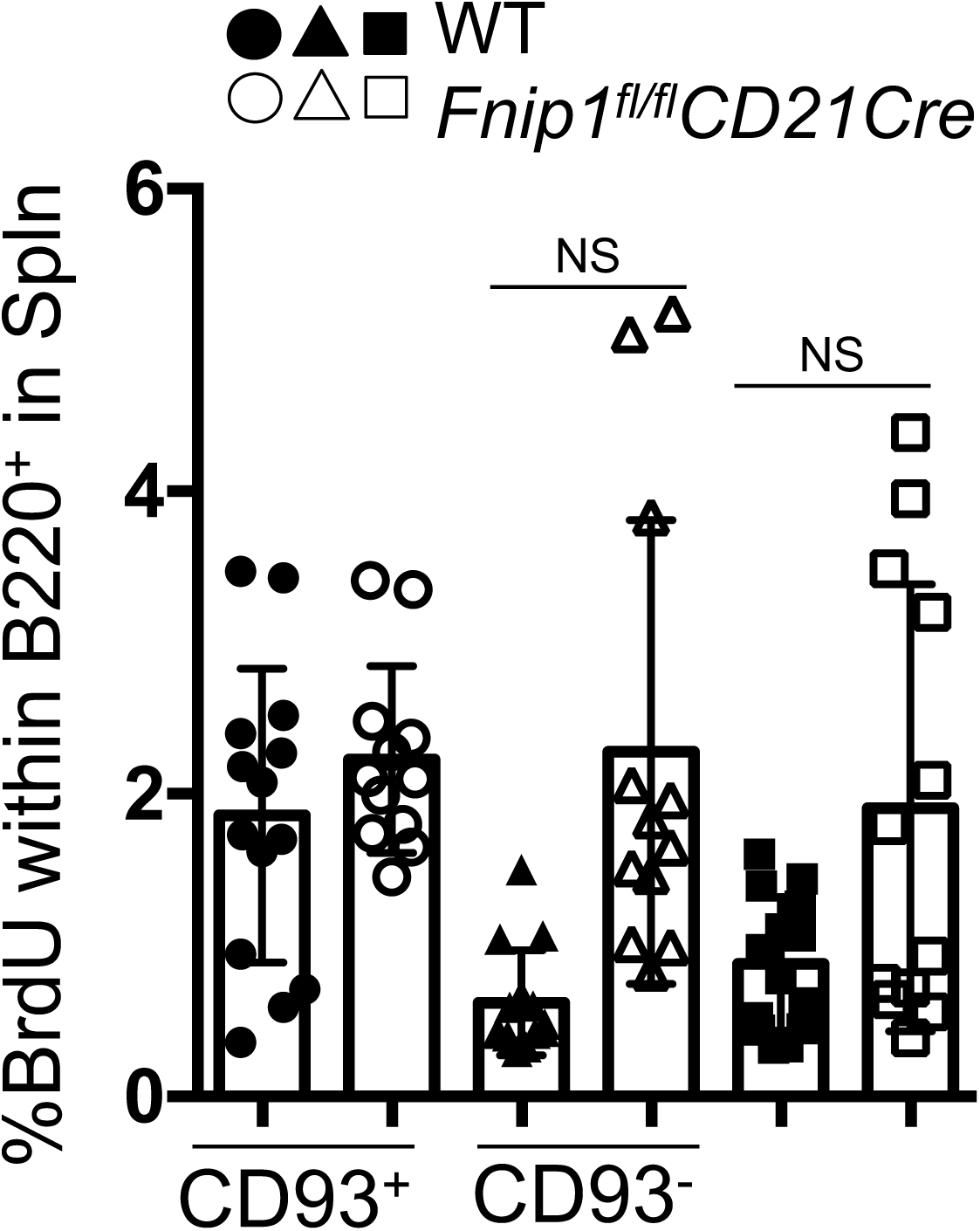

**Figure S9.**
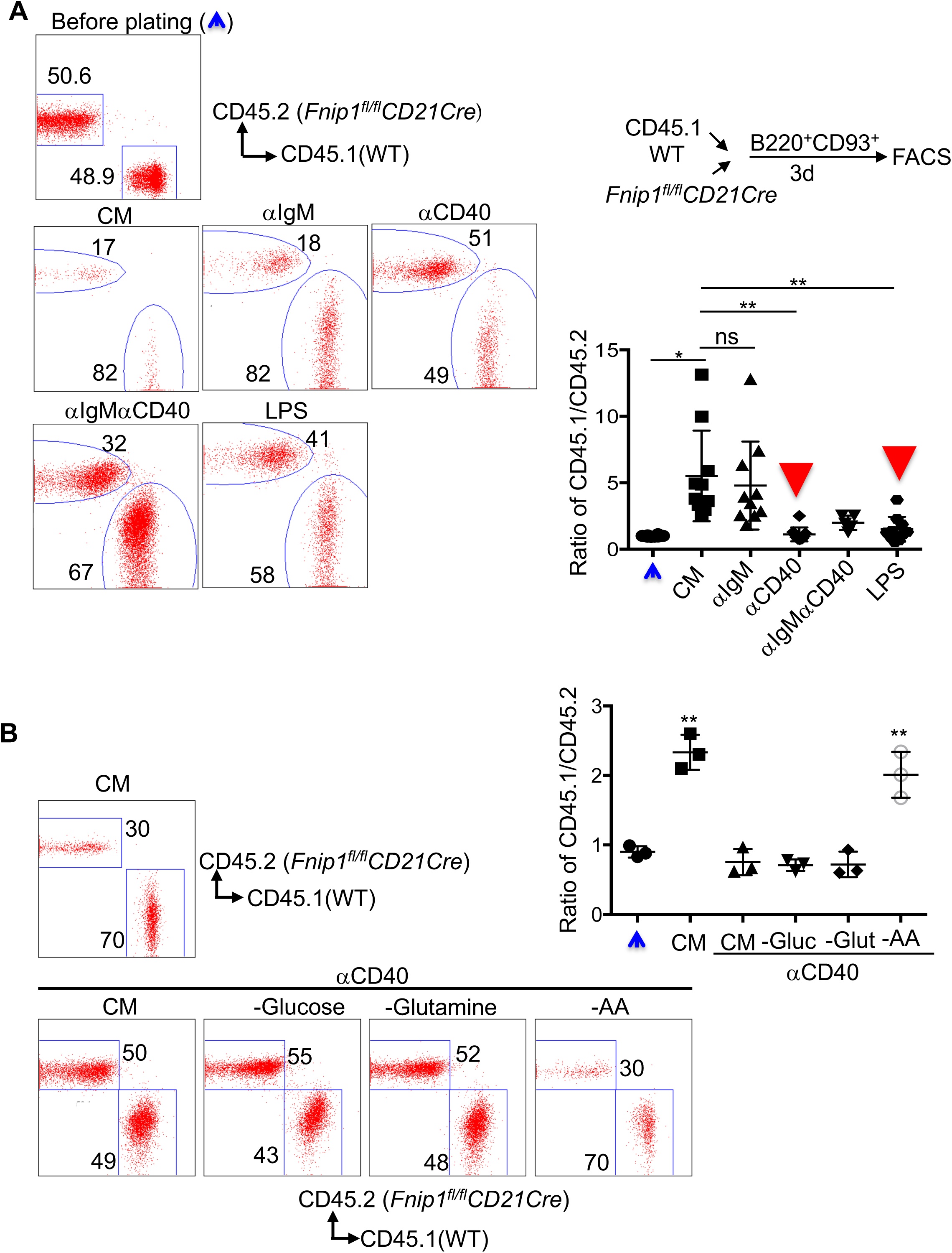

**Figure S10.**
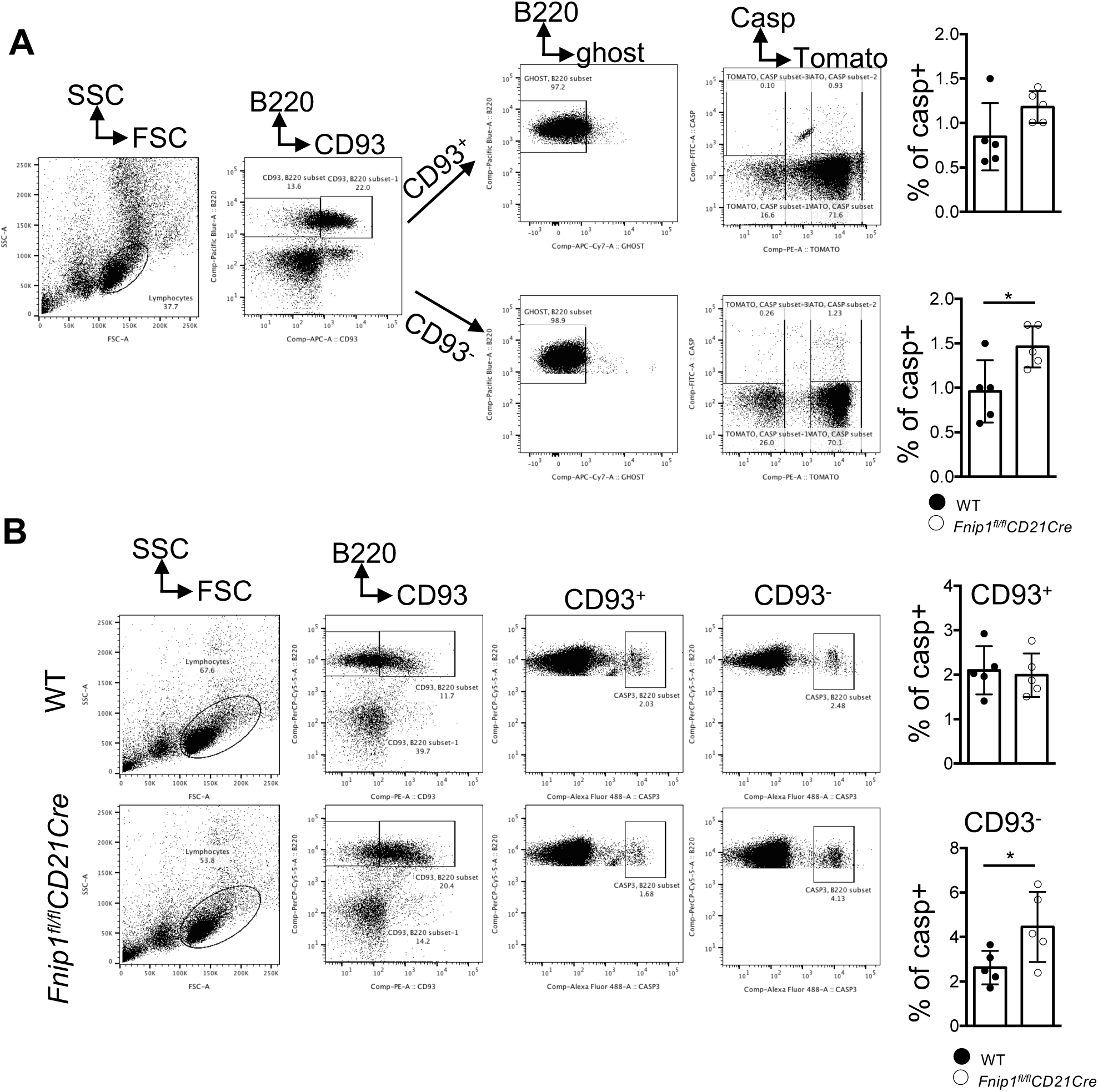

**Figure S11.**
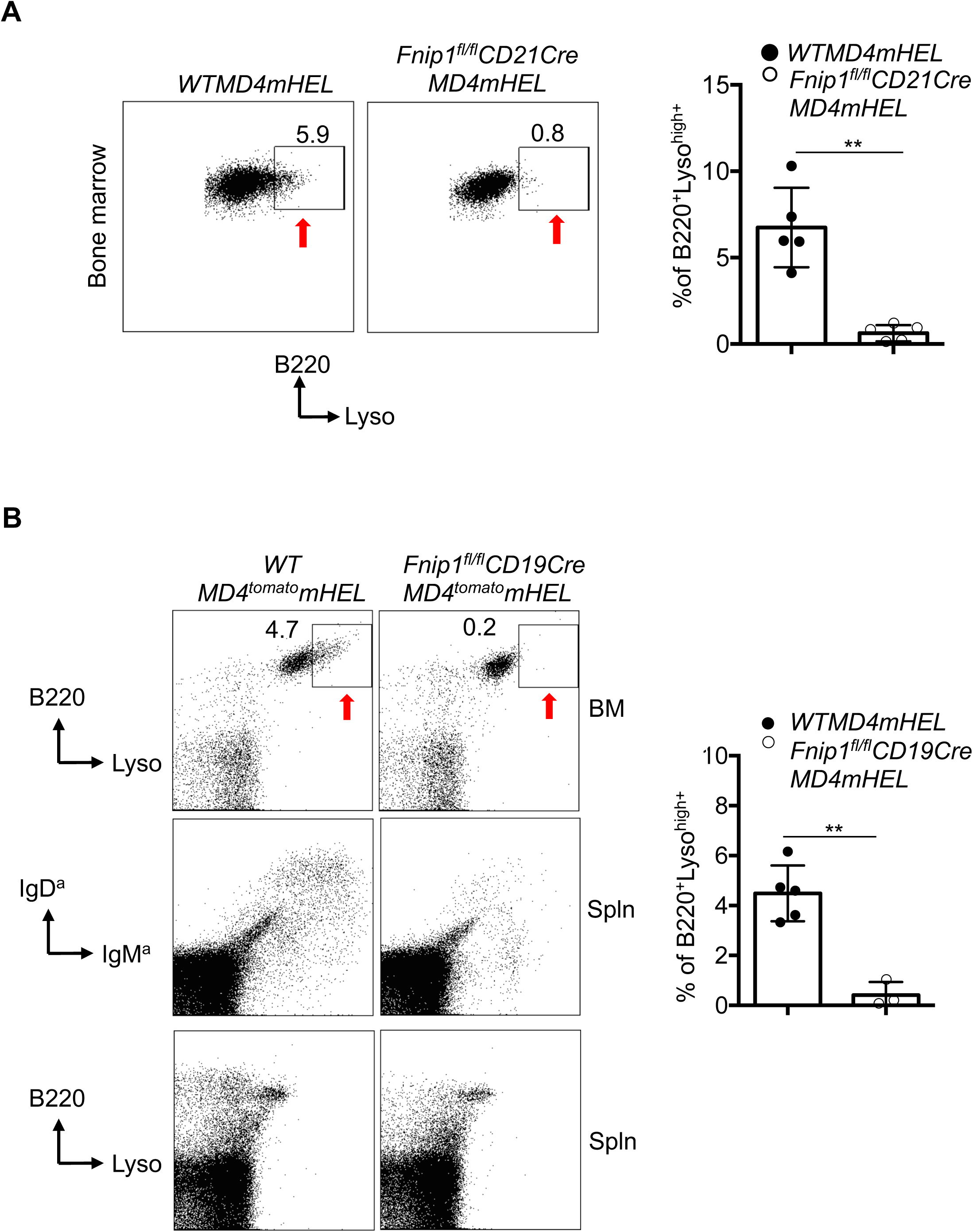

**Figure S12.**
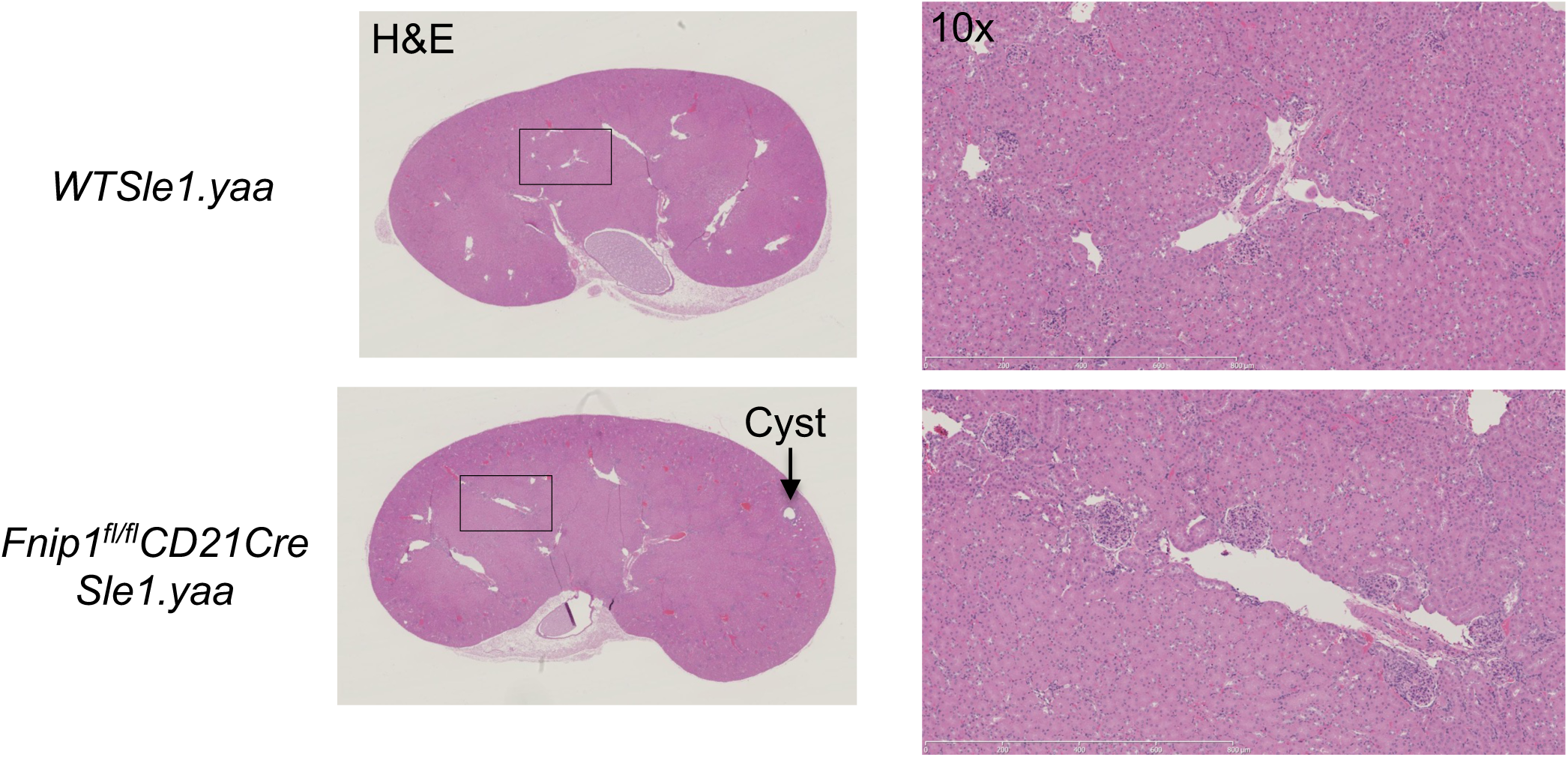

**Figure S13.**
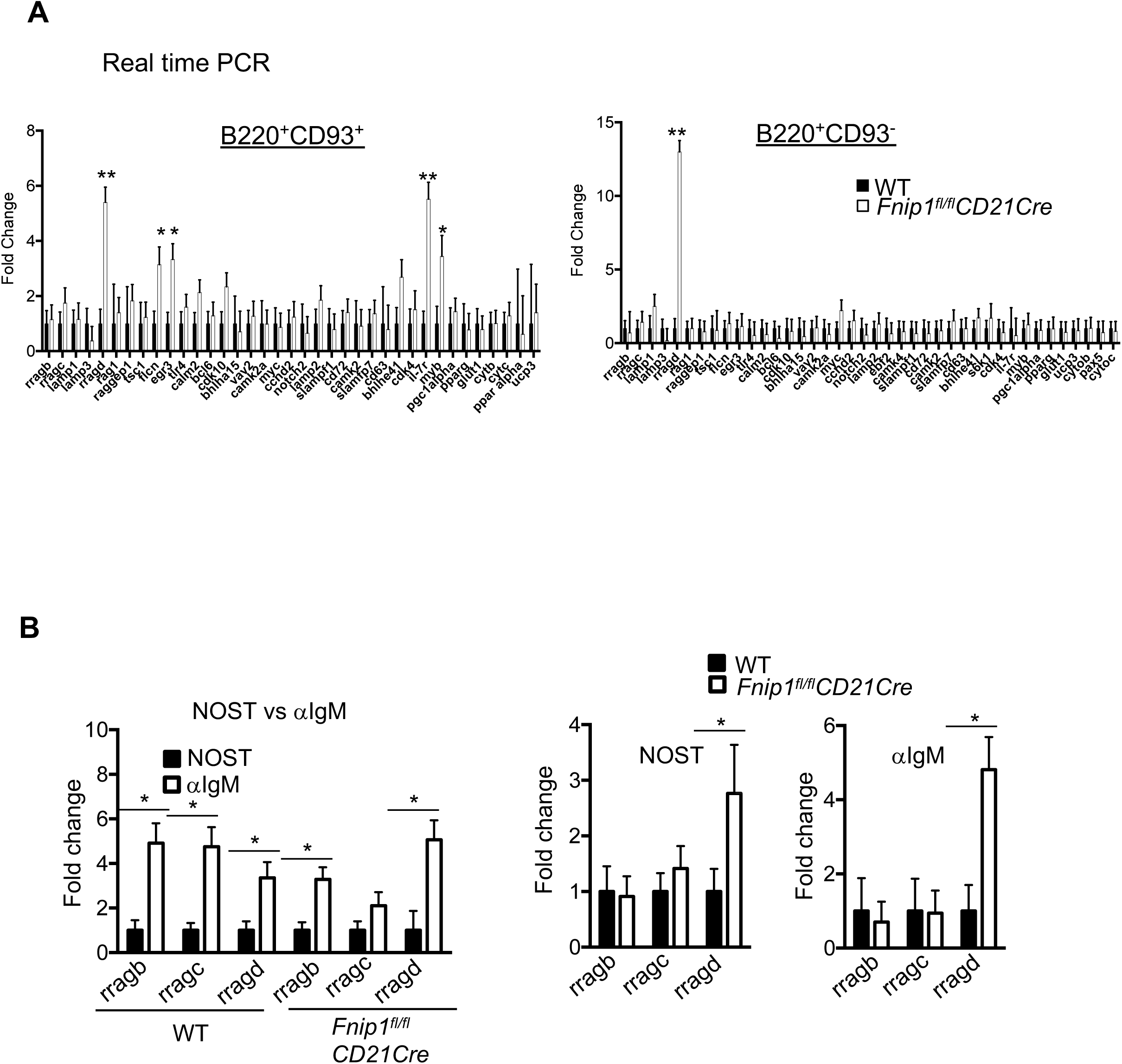

**Figure S14.**
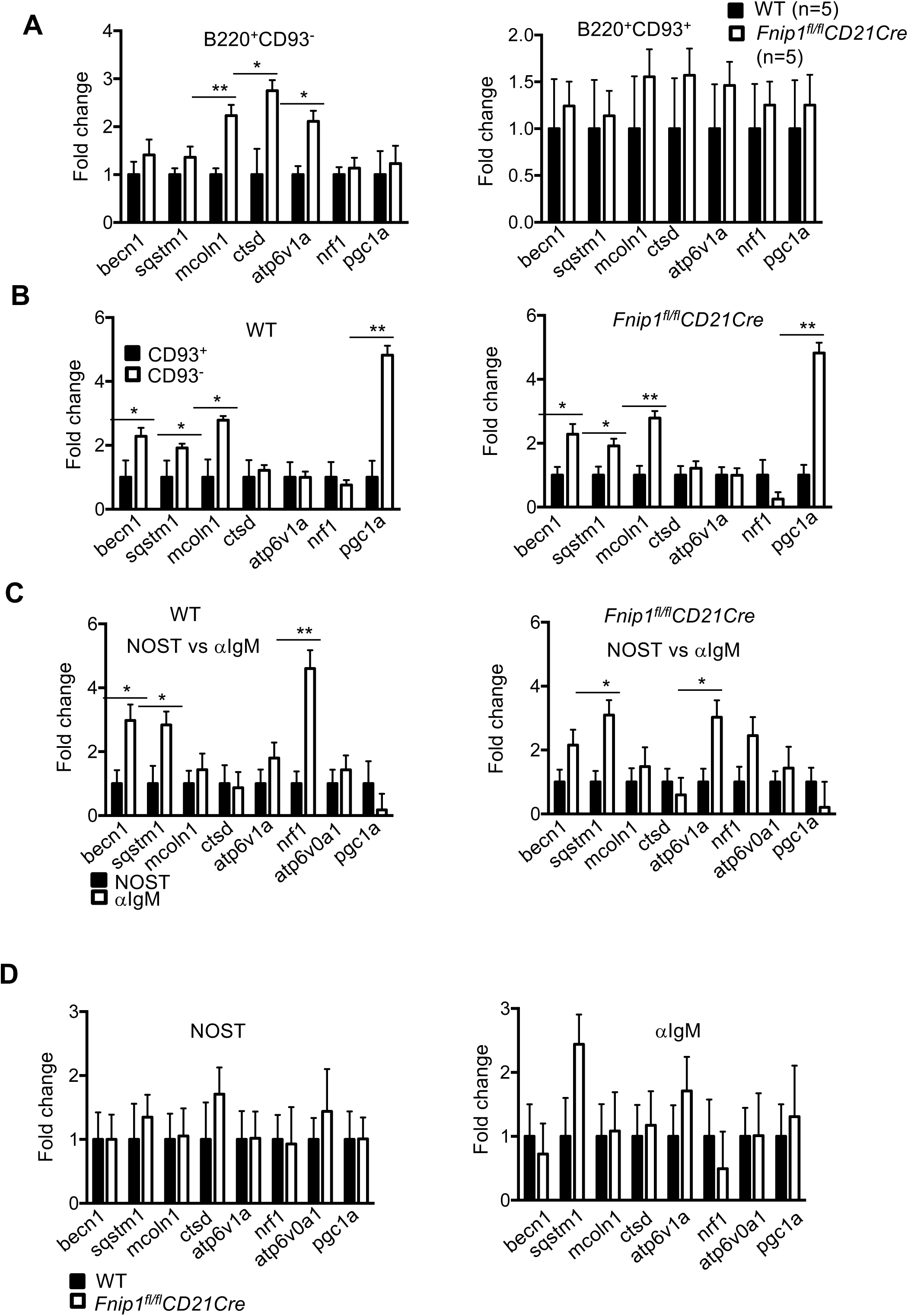

**Figure S15.**
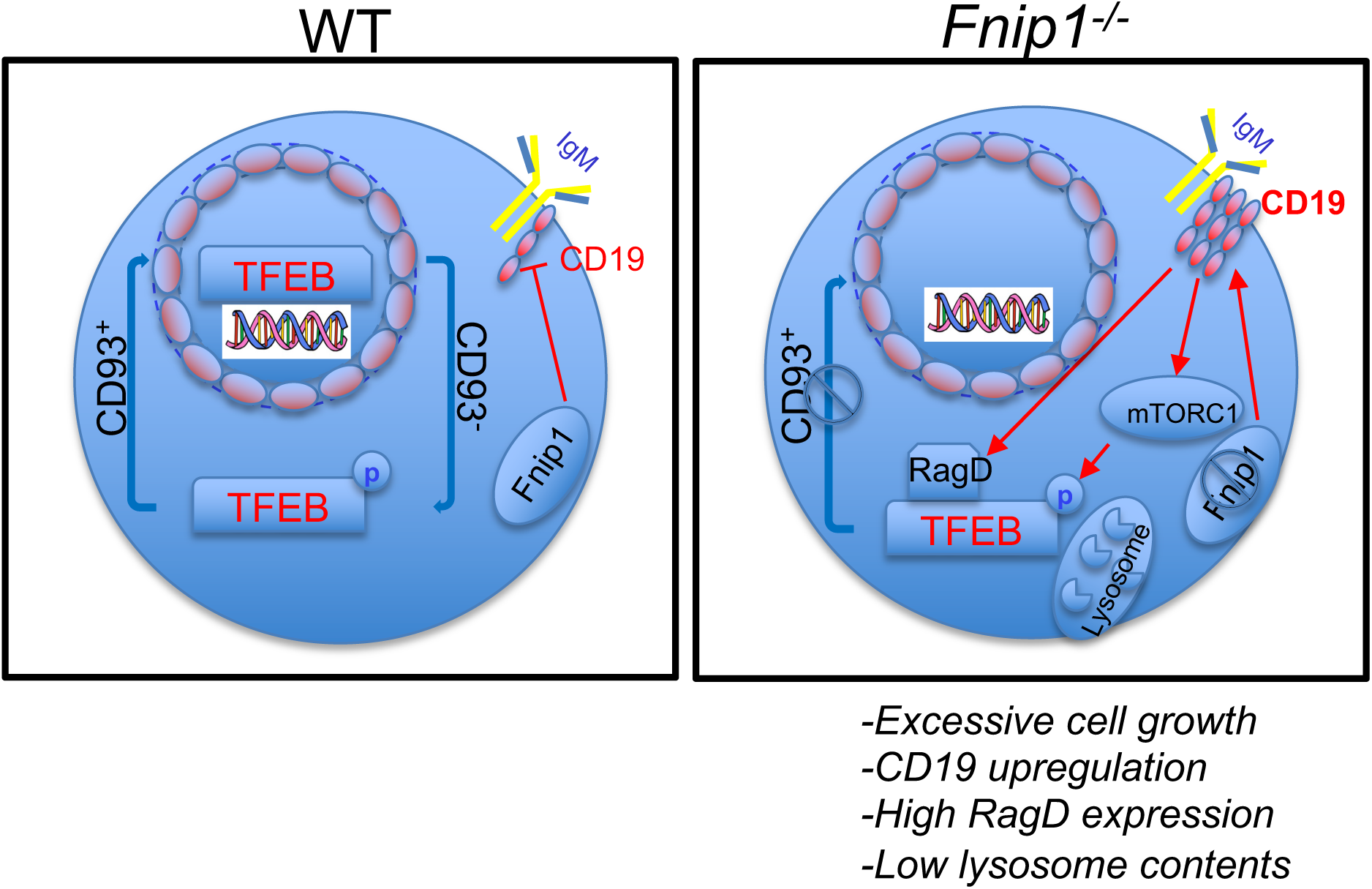
Proposed model of Fnip1 function

## Notes

### Competing Interest Statement

The authors have declared no competing interest.

