## Supplementary material for "FNIP1 Modulates B Cell Receptor Signaling Strength by Coordinating Metabolism During Development": NA

Supplemental Information

Figure S1 (related to Figure 1). Analysis of bone marrow, spleen, lymph node and peritoneal cavity shows a marked absence of mature B cells in *Fnip1*$^{fl/fl}$*CD21Cre* mice.

Figure S2 (related to Figure 2). *Fnip1*$^{fl/fl}$*CD21Cre* mice exhibit an increased frequency of pre-transitional (T0) B cells that retain the capacity to progress through the T1–T3 stages.

Figure S3 (related to Figure 2). Fnip1 deficiency in *Fnip1*$^{fl/fl}$*CD21Cre* mice impairs both T-dependent and T-independent immune responses, as well as germinal center B cell formation.

Figure S4 (related Figure 2). *Fnip1*$^{fl/fl}$*CD21Cre* mice maintain normal serum IgM levels and retain the capacity to generate plasmablasts.

Figure S5 (related Figure 2). *Fnip1*-deficient splenic B cells retain antigen-presenting capacity and effectively activate OTII $CD4^+$ T cells.

Figure S6 (related Figure 2). Fnip1 deficiency leads to a developmental arrest of splenic B cells at the transitional stage, accompanied by persistent Rag2 activity and elevated CD19 expression.

Figure S7 (related to Figure 3). Splenic B cells from *Fnip1*$^{fl/fl}$*CD21Cre* mice exhibit normal proliferative capacity.

Figure S8 (related to Figure 3). *Fnip1$^{fl/fl}$CD21Cre* mice exhibit normal BrdU incorporation.

Figure S9 (related to Figure 3 and Figure 7). Transitional B cells from *Fnip1$^{fl/fl}$CD21Cre* mice exhibit reduced viability in response to anti IgM stimulation, and their survival is dependent on amino acid availability.

Figure S10 (related to Figure 3). Fnip1 deficiency induces apoptosis during transitional B-cell development.

Figure S11 (related to Figure 4). Fnip1 deficiency leads to robust negative selection, resulting in the deletion of B cells expressing even low levels of surface IgM in response to membrane-bound HEL (mHEL).

Figure S12 (related to Figure 5). Fnip1 loss worsens polycystic kidney disease, causing enlarged cysts and increased glomerular diameter.

Figure S13 (related to Figure 7). RragD expression was selectively increased in *Fnip1*-deficient B cells, and this upregulation was dependent on BCR signaling strength.

Figure S14. TFEB-dependent gene expression is largely unaltered in Fnip1-deficient transitional B cells

Figure S15. Proposed model of Fnip1 function.

Supplemental Figure legends

Figure S1. *Fnip1$^{fl/fl}$CD21Cre* mice exhibit a marked loss of mature B cell populations across multiple lymphoid compartments.

(A) Bone marrow cellularity is comparable among WT, *Fnip1$^{-/-}$*, *Fnip1$^{fl/fl}$CD19Cre*, and *Fnip1$^{fl/fl}$CD21Cre* mice, as determined by manual cell counts using a hemocytometer. (B) Total splenocyte numbers in *Fnip1$^{fl/fl}$CD21Cre* mice are similar to those of WT controls. (C) *Fnip1$^{fl/fl}$CD21Cre* mice lack B220$^{+}$IgM$^{+}$ mature B cells in lymphoid tissues. (D) In the peritoneal cavity, B1a B cells are maintained at normal levels, whereas B2 and B1b B cell populations are markedly reduced in *Fnip1$^{fl/fl}$CD21Cre* mice. Cells were stained with fluorescent-conjugated antibodies against B220, CD5, and IgM. Data are presented as mean ± SD; **p<0.01.

Figure S2. *Fnip1$^{fl/fl}$CD21Cre* mice exhibit an increased frequency of pre-transitional (T0) B cells that retain the capacity to progress through the T1–T3 stages.

(A) Cells isolated from bone marrow, blood, and spleen were stained with fluorescent-conjugated antibodies against B220, CD93, IgM, and CD23. Pre-transitional B cells (T1-like; B220$^{+}$CD93$^{+}$IgM$^{low}$CD23$^{-}$) were identified by flow cytometry. (B) T0 cells from *Fnip1$^{fl/fl}$CD21Cre* mice differentiate into T1, T2, and T3 transitional B cells in vivo. A total of 1 × $10^6$ FACS sorted T0 cells were adoptively transferred into LPAB or *Rag2$^{-/-}$γc$^{-/-}$* recipient mice. Five to seven days post-transfer, splenic transitional B-cell populations were analyzed by flow cytometry using antibodies against B220, CD93, IgM, and CD23 to assess developmental progression. (C) To evaluate *Fnip1* deletion efficiency mediated by CD21Cre, FACS sorted T0, T1, T2, and T3 B cell populations were subjected to genomic PCR using a three-primer set (FW: CATCAGCTGTAAATGGCCTTC; FW: TGGTTCACAATGTGTGATGCATTT; RW: GTGGCTGTCCACTCCCTGATA). Tail DNA from the same mice was used as a control for floxed (fl) and deleted (d) alleles. Data are presented as mean ± SD; *p<0.05, **p<0.01.

Figure S3. Fnip1 deficiency impairs T-dependent and T-independent antibody responses and disrupts germinal center B cell formation in *Fnip1$^{fl/fl}$CD21Cre* mice.
(A and B) *Fnip1$^{fl/fl}$CD21Cre* (4-5 mice per group) and WT control mice (n=5) were immunized with either keyhole limpet hemocyanin (KLH; T-dependent antigen) or NP-Ficoll (T-independent antigen) as described in the Supplemental Methods. Serum was collected eight days post-immunization, and antigen-specific antibody isotypes were quantified by ELISA. Data are presented as mean ± SEM; *p<0.05, **p<0.01. (C and D) Mice were immunized with sheep red blood cells (SRBCs), and germinal center B cells were identified by flow cytometry using fluorescent-conjugated antibodies against B220, CD38, and CD95. Light zone and dark zone germinal center B cell subsets were distinguished using antibodies against CXCR4, and CD83. Data are presented as mean ± SD; **p<0.01.

Figure S4. *Fnip1$^{fl/fl}$CD21Cre* mice maintain normal serum IgM levels and retain the capacity to generate plasmablasts.
Positively selected B220$^{+}$ B cells ($1 \times 10^5$ cells) were cultured *in vitro* under plasmablast-inducing conditions with LPS (100 ng/mL) and IL-4 (50 ng/mL). Culture

supernatants were collected on day 4 from B cells stimulated with LPS alone, LPS plus IL-4, or anti-IgM (10μg/mL) /anti-CD40( 5 μg/mL). Cells were harvested on day 5.
(A and B) For total IgM quantification, culture supernatants (A) or serum samples (B) were analyzed by ELISA using IgM capture antibodies (Southern Biotech, cat# 1020-01) and IgM detection antibodies (SouthernBiotech, cat#. 1021-05) applied in serial dilutions. Colorimetric absorbance was measured at 405 nm. (C) Plasmablasts were identified as $B220^{low}CD138^{+}$ cells, with fluorochrome-conjugated antibodies, and analyzed by flow cytometry. The percentage of plasmablasts was quantified and compared between experimental groups (n = 4 per group). (D) FACS-sorted $IgD^{+}$ B cells ($1 \times 10^5$ cells) were stimulated *in vitro* with LPS and IL-4. Cells were harvested on day 4, fixed, permeabilized, and stained intracellularly with fluorochrome-conjugated anti IgG1 antibodies, followed by flow cytometric analysis. Data are presented as mean ± SD;*$p<0.05$, **$p<0.01$.

Figure S5. *Fnip1*-deficient splenic B cells retain antigen-presenting capacity and effectively activate OTII $CD4^{+}$ T cells.
(A and B) Positively selected $B220^{+}$ splenic B cells ($1x10^5$) from WT and *Fnip1*$^{fl/fl}$*CD21Cre* mice (n = 4 per group) were stimulated with LPS (100ng/ml) for 2 hrs, washed, and then co-cultured with CFSE-labeled OTII $CD4^{+}$ T cells ($1x10^5$ cells), which were isolated by positive CD4 selection, in the presence or absence of 20ug OVA peptide (GeneScript, Cat# RP10610-1). T cell activation was assessed 24 hours later by measuring CD25 and CD69 expression by flow cytometry, gating on $TCR\beta^{+}$ populations using fluorochrome-conjugated antibodies (A). Proliferation of OTII $CD4^{+}$ T cells was

evaluated 4 days post-culture by analyzing CFSE dilution (B). Data are presented as mean ± SD.

Figure S6. *Fnip1* deficiency leads to a developmental arrest of splenic B cells at the transitional stage, accompanied by persistent Rag2 activity and elevated CD19 expression.

(A) Splenic B cells from *WT Rag2GFP* and *Fnip1$^{fl/fl}$CD21CreRag2GFP* mice were stained with fluorochrome-conjugated antibodies against B220, CD93, IgM, IgD, and CD23 and analyzed by flow cytometry. The pre-transitional B cells in T0 compartment was defined as $B220^{+}IgM^{low}IgD^{hi}CD23^{-}$. (B) Rag2 expression across stage-specific splenic B cell subsets is shown. Splenic cells from *WT Rag2GFP* and *Fnip1$^{fl/fl}$CD21CreRag2GFP* mice were stained and analyzed by flow cytometry. B cells were gated based on CD19, IgM, CD93, CD23, and Rag2GFP. *Fnip1$^{fl/fl}$CD21Cre* mice exhibited a block at the transitional B cell stage, as indicated by an increased frequency of $B220^{+}CD93^{+}$ cells. *Fnip1*-deficient splenic B cells displayed a distinct $CD19^{hi}Rag2^{-}CD23^{-}IgM^{high/low}$ phenotype with excessive cell growth corresponding to the T0 and T1 compartments prior to maturation. Data are presented as mean ± SD; *p<0.05, **p<0.01.

Figure S7. Splenic B cells from *Fnip1$^{fl/fl}$CD21Cre* mice exhibit normal proliferative capacity.

(A-C) Purified splenic B cells (1 × $10^5$) from *Fnip1$^{fl/fl}$CD21Cre* mice were labeled with CFSE and stimulated in vitro with (A) anti IgM, (B) LPS, or (C) anti CD40 antibodies. Proliferation was assessed by CFSE dilution using flow cytometry. (D) Splenic B cells were isolated from CD45.1$^+$ WT and CD45.2$^+$ *Fnip1$^{fl/fl}$CD21Cre* mice, labeled with CFSE, and co-transferred in equal numbers (5 × $10^6$ cells each) into CD45.2$^+$ LPAB recipient mice. Anti IgM was administered intraperitoneally 24 hours post-transfer. CFSE dilution was analyzed by flow cytometry four days later to assess in vivo B cell proliferation. Data are presented as mean ± SD.

Figure S8. BrdU incorporation is preserved in *Fnip1$^{fl/fl}$CD21Cre* mice.
BrdU incorporation was evaluated by flow cytometry using fluorescent-conjugated antibodies against B220, CD93, and BrdU. The percentage of BrdU$^+$ cells was quantified within the B220$^+$CD93$^+$ (immature) and B220$^+$CD93$^-$ (mature) splenic B cell populations. Data are presented as mean ± SD; ns means not significant.

Figure S9. Transitional B cells from *Fnip1$^{fl/fl}$CD21Cre* mice exhibit impaired survival upon anti IgM stimulation and are sensitive to amino acid availability.
(A) CD93$^+$ splenic B cells were isolated from CD45.1$^+$ WT and CD45.2$^+$ *Fnip1$^{fl/fl}$CD21Cre* mice. Equal numbers of cells (2 × $10^5$ total) were mixed at a 1:1 ratio and co-cultured under various stimulatory conditions (anti IgM, anti CD40, anti IgM plus anti CD40, or LPS). After 72 hours, the ratio of CD45.1 to CD45.2 cells was assessed by flow cytometry to evaluate relative survival. (B) CD93$^+$ splenic B cells (B220$^+$CD93$^+$) from WT (CD45.1) and *Fnip1$^{fl/fl}$CD21Cre* (CD45.2) mice were co-cultured (1:1 ratio; 2 × $10^5$ total cells) in 96-well flat-bottom plates under media conditions with or without glucose, glutamine, or amino acids, in the presence of anti CD40 (5 μg/mL). Cell survival was assessed after 72 hours by flow cytometry. Data are presented as mean ± SD; *p<0.05, **p<0.01.

Figure S10. Fnip1 deficiency induces apoptosis during transitional B-cell development.
(A) Splenic cells from *Fnip1fl/flCD19CreTomato* mice (n = 5) were stained with fluorochrome-conjugated antibodies and analyzed by flow cytometry. B cells were gated based on B220, CD93, a viability dye (Ghost), and Tomato expression. Tomato negative cells (wild-type *Fnip1*) and Tomato positive cells (*Fnip1* deleted) were analyzed for the percentage of caspase 3/7 positive cells among $CD93^+$ and $CD93^-$ B-cell populations.
(B) Similarly, splenic cells from *Fnip1*$^{fl/fl}$*CD21Cre* mice and WT controls (n = 5) were stained and analyzed by flow cytometry. Data are presented as mean ± SD; statistical significance was assessed using an unpaired two-tailed Student's *t* test. *p<0.05.

Figure S11. Fnip1 deficiency leads to robust negative selection, resulting in the deletion of B cells expressing even low levels of surface IgM in response to membrane-bound HEL (mHEL).
(A) Bone marrow cells from *WTMD4mHEL* and *Fnip1*$^{fl/fl}$*CD21CreMD4mHEL* mice were stained with fluorescent-conjugated antibodies against B220, and lysozyme, and analyzed by flow cytometry. The frequency of $B220^+Lyso^{high}$ B cells was quantified.
(B) Bone marrow and spleen cells from *WTMD4mHEL* and *Fnip1*$^{fl/fl}$*CD19CreMD4mHEL* mice were stained with antibodies against B220, $IgD^a$, $IgM^a$, and Lysozyme and analyzed by flow cytometry to assess the extent of B cell deletion. Data are presented as mean ± SD; **p<0.01.

Figure S12. Fnip1 loss worsens polycystic kidney disease, causing enlarged cysts and increased glomerular diameter.

Kidneys from 3-month-old *WTSle1.yaa* and *Fnip1$^{fl/fl}$CD21CreSle1.yaa* mice were stained with hematoxylin and eosin (H&E) at the Histology and Imaging Core (HIC) at the University of Washington.

Figure S13. *RragD* expression is increased in CD93$^{low}$ cells from *Fnip1$^{fl/fl}$CD21Cre* mice, and this upregulation is dependent on BCR signaling strength.
(A) cDNA was synthesized from FACS-sorted B220$^{+}$CD93$^{+}$ and B220$^{+}$CD93$^{-}$ splenic B cells (n=5) and analyzed by quantitative real-time PCR using gene-specific primers listed in supplemental methods.  (B) Positively selected B220$^{+}$ splenic B cells were stimulated with or without anti-IgM antibodies for 6 h, followed by cDNA synthesis and quantitative real-time PCR analysis of autophagy-related genes, lysosomal genes, mitochondrial genes, and *Rrag* family members (*Rragb, Rragc,* and *Rragd*) (n = 4). Relative gene expression levels were compared between genotypes under basal and stimulated conditions. Data are presented as mean ± SEM; *p<0.05, **p<0.01.

Figure S14. TFEB-dependent gene expression is largely unaltered in Fnip1-deficient transitional B cells.
(A and B) cDNA was generated from FACS-sorted B220$^{+}$CD93$^{+}$ and B220$^{+}$CD93$^{-}$ splenic B cell populations (n = 5 per genotype) and analyzed by quantitative real-time PCR using gene-specific primers targeting autophagy-related genes (*Becn1, Sqstm1*), lysosomal genes (*Mcoln1, Ctsd*), and mitochondrial genes (*Atp6v1a, Nrf1, Pgc1a*), as listed in the Supplemental Methods. Gene expression levels were compared between WT and *Fnip1$^{fl/fl}$CD21Cre* B cells within the CD93$^{+}$ and CD93$^{-}$ populations. (C and D) Positively selected B220$^{+}$ splenic B cells were stimulated with or without anti-IgM antibodies for 6 h, followed by cDNA synthesis and quantitative real-time PCR analysis

of autophagy-related genes, lysosomal genes, and mitochondrial genes. Relative gene expression levels were compared between genotypes under basal and stimulated conditions. Data are presented as mean ± SEM; *$p<0.05$, **$p<0.01$.

Figure S15. Model summarizing the proposed function of Fnip1 during the maturation of B cells.

Fnip1 acts as a critical sensor that integrates B cell receptor (BCR) signaling strength with metabolic state of developing B cells. In the absence of Fnip1, altered BCR signaling thresholds and enhanced CD19-dependent signaling lead to increased RragD expression and activation of mTORC1. As a result, Fnip1-deficient B cells exhibit excessive cell growth, upregulation of CD19, elevated RagD levels, and reduced lysosomal content.

### Supplemental Experimental Methods

*Immunization*

Mice were immunized intraperitoneally with KLH (100ug) emulsified in Imject Alum Adjuvant, NP-Ficoll (50 µg) or sheep red blood cells (SRBCs; 100µl in PBS). SRBCs were washed twice in PBS before injection. Serum was collected by cardiac puncture 8 days post-immunization.

*In Vitro and in Vivo proliferation assays*

CFSE-labeled splenic B cells were stimulated with anti IgM (10 µg/mL), anti CD40 (5 µg/mL), or LPS (100 ng/mL) for 72 hours in vitro. Proliferation was assessed by CFSE dilution within B220$^+$ cells. For in vivo proliferation, WT (CD45.1) and *Fnip1$^{fl/fl}$CD21Cre* (CD45.2) B cells were CFSE-labeled and co-transferred into LPAB (Fnip1$^{-/-}$) mice. Anti IgM (150 µg) was injected intraperitoneally 24 hours later. Proliferation was assessed three days post-stimulation.

*Nutrient deprivation assays*

Glucose-free or glutamine-free RPMI media were supplemented with 10% dialyzed fetal calf serum. Amino acid-restricted media were formulated based on RPMI1640 composition and adjusted to pH7.2.  HBSS solution was supplemented with MEM vitamins, non-essential amino acids, L-glutamine (2mM), sodium pyruvate (1mM), glucose (11mM), 10% dialyzed FCS, supplemented with or without essential amino acids.  Transitional splenic B cells (B220$^+$CD93$^+$) from WT (CD45.1) and *Fnip1$^{fl/fl}$CD21Cre* (CD45.2) mice were isolated using B cell isolation kit /CD93 microbeads and then co-cultured in a 1:1 ratio (2x10$^5$ total cells) in 96-well flat-bottom plates, with or without anti-CD40 (5 µg/mL). After 72 hours, survival was assessed by flow cytometry.

*H&E staining and glomerular diameter measurement*

Kidneys were fixed in 10% formalin and processed by the Histology and Imaging Core (HIC) at the University of Washington (Seattle) for hematoxylin and eosin (H\&E) staining. Slides were digitally scanned at the HIC, and glomerular diameters were measured using NDP.view2 software at 40x magnification. The longest diameter of each glomerulus was recorded for analysis.

*Primers used for real-time PCR*

Camk2a FW:ACCCTCTACTTTCTCTCCTCC 12322(gene ID) RW:ACTTTGGTGTCTTCGTCCTC; CD63 FW:TTTGCTCTACGTTCTCCTGC 12512 RW:AAGACAACCTGAACCGCTAC; Lamp1 FW:ACAAACCCCACTGTATCCAAG 16783 RW:CATTTGGGCTGATGTTGAACG; Lamp2 FW:GGTATTCACCTGCAAGCTTTTG 16784 RW:GGGTGTTGAAGTTGGAGTGAG; Lamp3 FW:TAGCACTGATTGTTCAAGAAAAGG 239739 RW:GATATTCACAGATCCACCCTGG; rragB FW:ACAGAGAAAGAGAATGTGGGC 245670 RW:ACCAGATCCACTTTTACCCATC; rragC FW:AGCTCCTTTGTGAACTTCCAG 54170 RW:GGTTAAAGCCTCCATGTAGTCG; rragD FW:TTTGACCCTACCTTTGACTATGAG 52187 RW:ATCAGTGTTCACCTTGTAGGC; Rag1 FW:ACCTGAAGATGAAACCCGTG 19373 RW:GTAATTGGTGATTTTGCCCTCG; S6k1 FW:ACAGGAGCAAATACTGGGAAG 72508 RW:TCAGGTCCACAATGAAAGGG; Myc FW:GCTGTTTGAAGGCTGGATTTC 17869 RW:GATGAAATACTGTACGGAG; Cdk4 FW:ACAAGTAATGGGACCGTCAAG 12567 RW:GGGTGTTGCGTATGTAGACTG; Ccnd2 FW:GTGTTCCTATTTCAAGTGCGTG 12444 RW:AGCCAAGAAACGGTCCAG; Tlr4 FW:TTCAGAACTTCAGTGGCTGG 21898 RW:TGTTAGTCCAGAGAAACTTCCTG; CD72 FW:CATCTCACATACCCTCAGAAGTC 12517 RW:TTCGAACGATCTAGCAACTCC; Vav2 FW:ATCAGGCCATTTCCATCAGAG 22325 RW:TTCATCTTCACACGGGACAC; Calm2 FW:GTGTTTGATAAGGATGGCAATGG 12314 RW:CTCTTCGTAGTTTACCTGACCG; Camk4 FW:CAGGATTGTGGAGAAGGGATAC 12326 RW:TCTGGTTTGAGGTCACGATG; Bcl6 FW:GGAAACCCAGTCAGAGTATTCG 12053 RW:GGCAGCGATCACATTTGTATG; Tsc1 FW: TCTACTTCCACCCCTTCCTC 64930 RW:CATACCACAGACCGCAGATG; Myb FW:CAAGGGAAGAGGATGAGAAGC 17863 RW:ATGAGTTCAGGGTTCAGCAC; IL-7R FW:TCTGGAGAAAGTGGAAATGCC 16197 RW:AGCTGTGTTGATGTCTGAGTC; Camk2b FW:TCAAGCCCCAGACAAACAG 12323 RW:ATTCCTTAATCCCGTCCACTG; Slamf1 FW:TCTCCCTGGCTTTTGAGTTG 27218 RW:GATGCGGACACTTTTGTTCAC; Ebf2 FW:TCTCCAGGTTGTGTTTGGTAC 13592 RW:GCTCCTTTGCAGAACTGTTTG; Notch2 FW:AAAATCTGCCCTCCACTGG 18129 RW:CCGCTTCATAACTTCCCTCTC; Bhlhe41 FW:GTGTAAACCCAAAAGGAGCTTG 79362 RW:CAATTTCAGATGTTCGGGCAG; Flcn FW:CTATCCCTGCCCATGTTCTG 216805 RW:GTGGAGGTTTGAGCGAGTG; Egr3 FW:TCGGTAGCCCATTACAATCAG 13655 RW:AGCGAACTTTCCCAAGTAGG; Cdk10 FW:CGGATGGACAAAGAGAAGGATG 234854 RW:CAGTAACCCATGACCAGGAAG; Rapgef1 FW:TCATCTCCAAAATGAAGCCCC 107746 RW:ATTGTCCTCACTTTCTCGGC; Slamf7 FW:ATGACACAATCCCTTACACGG 75345 RW:AGCTTAATGACCTTGGCACG; Bhlha15 FW:CCAGGCCCTAAATTATACCAGC 17341 RW:CTGTGTAGAGTAGCGTTGCAG; Ppar alpha FW:TCGGCGAACTATTCGGCTG

19013 RW:GCACTTGTGAAAACGGCAGT; Ppargamma FW:TGTGGGGATAAAGCATCAGGC 19016 RW:CCGGCAGTTAAGATCACACCTAT; Ucp3 FW:GAGATGGTGACCTACGACATCA 22229 RW:GCGTTCATGTATCGGGTCTTTA; Pax5 FW:AGTGTCTACAGGCTCCGTGA 18507 RW:CCCTCTTGCGTTTGTTGGTG; Pgc1 alpha FW:AGCCAAACCAACAACTTTATCTCTTC 19017 RW:TTAAGGTTCGCTCAATAGTCTTGTTC; Glut1 FW:TCAACGAGCATCTTCGAGAAGGCA 20525 RW:TCGTCCAGCTCGCTCTACAACAAA; CytoB FW:AAAGCCACCTTGACCCGATT 17711 RW:GATTCGTAGGGCCGCGATA; Mt-Co2 FW:CCATAGGGCACCAATGATACTG 17709 RW:AGTCGGCCTGGGATGGCATC; Nrf1 FW: GGCAACAGTAGCCACATTGGCT 18181 RW: GTCTGGATGGTCATTTCACCGC; CTSD FW: TAAGACCACGGAGCCAGTGTCA 13033 RW: CCACAGGTTAGAGGAGCCAGTA; Mcoln1 FW: GTCGGTGTCATTCGCTACCTGA 94178 RW: GAACGATCCAGCCACAGAAGCA; Becn1 FW: CAGCCTCTGAAACTGGACACGA 56208 RW: CTCTCCTGAGTTAGCCTCTTCC; Sqstm1 FW: GCTCTTCGGAAGTCAGCAAACC 18412 RW: GCAGTTTCCCGACTCCATCTGT; Atp6v1a FW: GCTGGCTTCTTTCTATGAGCGAG 11964 RW: GCGTTGCAGAAGTGACTGGATC; Atp6v0a1 FW: CTGTTATCCTCGGCATCATCCAC 11975 RW: CAGGTAGCCAAACAACGAGGAC.
